# Functional substitutions of amino acids that differ between GDF11 and GDF8 impact skeletal development and skeletal muscle

**DOI:** 10.1101/2022.05.09.491247

**Authors:** John Lian, Ryan G. Walker, Andrea D’Amico, Ana Vujic, Melanie J. Mills, Kathleen A. Messemer, Kourtney R. Mendello, Jill M. Goldstein, Krystynne A. Leacock, Soraya Epp, Emma V. Stimpfl, Thomas B. Thompson, Amy J. Wagers, Richard T. Lee

## Abstract

Growth differentiation factor 11 (GDF11) and GDF8 (MSTN) are closely related TGFβ family proteins that interact with nearly identical signaling receptors and antagonist proteins. However, GDF11 appears to activate SMAD2/3 *in vitro* and *in vivo* more potently than GDF8. The ligands possess divergent structural properties, whereby substituting unique GDF11 amino acids into GDF8 enhanced activity of the resulting chimeric GDF8. We investigated potentially distinct endogenous activities of GDF11 and GDF8 *in vivo* by genetically modifying their mature signaling domains. Full recoding of the GDF8 mature domain to that of GDF11 yielded mice lacking GDF8, with GDF11 levels ∼50-fold higher than normal, and exhibiting modestly decreased muscle mass, with no apparent negative impacts on health or survival to adulthood. Substitution of two specific amino acids in the fingertip region of GDF11 with the corresponding GDF8 residues resulted in prenatal axial skeletal transformations, consistent with *Gdf11*-deficient mice, without apparent perturbation of skeletal or cardiac muscle development or homeostasis. These experiments uncover distinctive features between the GDF11 and GDF8 mature domains *in vivo* and identify specific requirement for GDF11 in early-stage skeletal development.

**Summary Statement:** Replacement of amino acids unique to GDF11 and GDF8 into the native locus of the other ligand yields measurable, differential skeletal and muscle phenotypes, revealing distinct features between the ligands.

## Introduction

The transforming growth factor beta (TGFβ) superfamily of proteins is well-known for regulating embryological development, wound healing, and adult tissue maintenance. In recent years, two highly homologous TGFβ proteins—Growth Differentiation Factor 11 (GDF11) and GDF8 (also known as myostatin/MSTN)—have garnered substantial interest with evidence of their roles in aging and regenerative processes (Walker et al., 2016; Loffredo et al., 2013; Du et al., 2017; Poggioli, 2015; Biesemann et al., 2014; Sinha et al., 2014). Due to the 89% amino acid sequence identity in their C-terminal signaling domains, GDF11 and GDF8 have been viewed as serving redundant functions *in vivo* (Poggioli et al., 2015; Walker et al., 2016; McPherron et al., 2009) (**Fig. 1A**). Yet, growing evidence suggests that GDF11 and GDF8 have distinct potencies and different spatiotemporal functions *in vivo*. As members of the activin subclass of TGFβ, both GDF8 and GDF11 signal through the type I receptors ALK4, ALK5 and ALK7 (Walker et al., 2017; Rebbapragada et al., 2003; Andersson et al., 2006). Molecularly, they are synthesized as precursors that remain in an inactive, latent complex until a Tolloid-like (TLD) protease cleaves the ligand prodomain to relieve the mature domain from inhibition (Rebbapragada et al., 2003; Wolfman et al., 2003; Ge et al., 2005; McFarlane et al., 2005; Lee & McPherron, 2001; Anderson et al., 2008). Mature GDF11 and mature GDF8 each consist of two monomers linked by disulfide bonds to form a homodimer of propeller-like shape (**Fig. 1B**), which creates symmetrical concave and convex surfaces used for receptor binding (Walker et al., 2017; Yadin et al., 2016). To signal, the ligands assemble a combination of two type II and two type I Ser/Thr kinase receptors that have a single extracellular ligand-binding domain (Allendorph et al., 2006; Weber et al., 2007). Assembly of this complex allows the type II receptor to phosphorylate the type I receptor, which initiates the SMAD signaling cascade (Weiss & Attisano, 2013).

**Figure 1.**
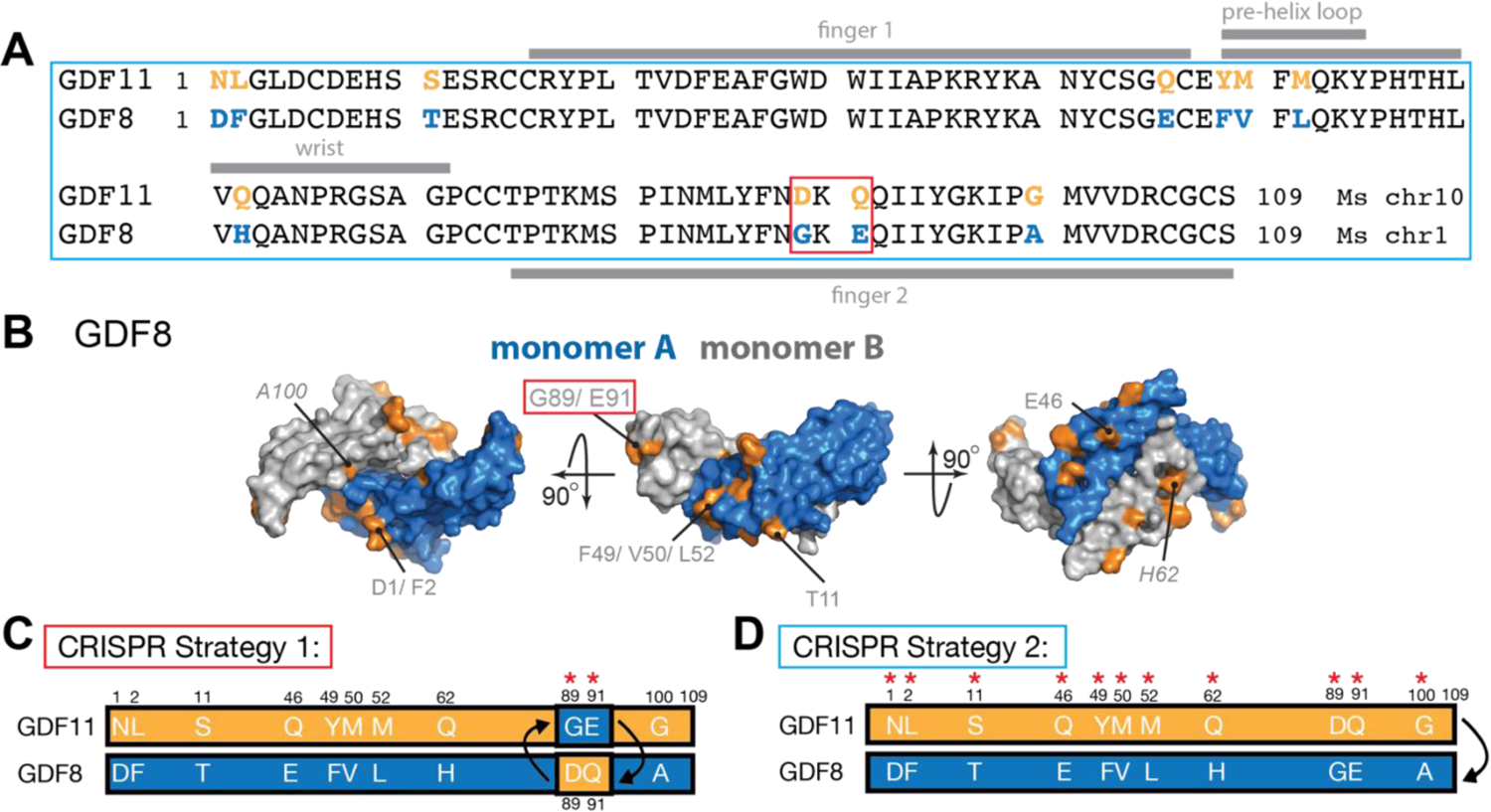
Sequence differences between *Gdf11* and *Gdf8* and CRISPR/Cas9 strategy to substitute amino acid residues within GDF11 and GDF8 mature domains. **(A)** *Gdf11* unique amino acid residues are highlighted in orange, and *Gdf8* unique residues are in blue. **(B)** Surface representation of GDF8 (monomer A in blue, monomer B in grey) showing location of unique GDF8 amino acid residues (in orange), with G89 and E91 highlighted. **(C)** Schematic of CRISPR/Cas9 strategies for changing residues in *Gdf11* and *Gdf8* native loci. In Strategy 1, the D89 and Q91 amino acid residues within *Gdf11* are changed for the analogous G89 and E91 amino acid residues from *Gdf8*, and the G89 and E91 amino acid residues within *Gdf8* are changed for the analogous D89 and Q91 residues from *Gdf11*. **(D)** In Strategy 2, the full mature domain of *Gdf8* is changed for the full mature domain from *Gdf11*. Amino acid residues shown for *Gdf11* (orange) and *Gdf8* (blue).

GDF8 is expressed postnatally, predominantly by skeletal and cardiac muscles, and is a well-recognized negative regulator of muscle growth (McPherron et al., 1997; McPherron & Lee, 1997). GDF11 is expressed more broadly across multiple tissues, with noted involvement in early development (McPherron et al., 1999; McPherron et al., 2009; Liu, 2006; Wu et al., 2003; Kim et al., 2005). Genetic depletion of *Mstn* (*Gdf8*) results in hyper-muscular, hypo-adipose phenotypes in numerous animal species, including humans (McPherron et al., 1997; McPherron & Lee, 1997; Schuelke et al., 2004; Mosher et al., 2007; Clop et al., 2006), whereas homozygous deletion of *Gdf11* leads to axial skeletal malformation and defects in organ development in mice (McPherron et al., 1999). In addition, recent evidence suggests that genetic loss of GDF11 function in humans causes multi-system pathology with variable impact on the skeleton, nervous system, heart, muscle and/or connective tissue (Ravenscroft et al., 2021). Importantly, *Gdf11*-null mice exhibit perinatal lethality, whereas *Mstn*-null (*Gdf8^-/-^*) mice do not (McPherron et al., 1999), and lower levels of *Mstn* in *Gdf8^+/-^* heterozygotes may actually extend lifespan (Mendias et al., 2015). These differences in postnatal survival following genetic manipulation make comparative studies of *Gdf11* versus *Gdf8* activities *in vivo* particularly difficult, while reports on GDF11’s essential functions in adulthood continue to be inconsistent and controversial. Nevertheless, interest persists from pharmaceutical and biotechnology companies in the potential effects of GDF11 in age-related organ dysfunction (Loffredo et al., 2013; Katsimpardi et al., 2014; Sinha et al., 2014), and several studies support the notion that exogenous GDF11 may regulate cardiac hypertrophy and skeletal muscle repair in older animals (Loffredo et al., 2013; Du et al., 2017; Sinha et al., 2014).

In a prior study, we demonstrated that GDF11 and GDF8 differ in their signaling properties in multiple cell lines and cultured primary myoblasts, with GDF11 signaling much more potently than GDF8 and more efficiently utilizing the type I receptors ALK4, ALK5, and ALK7 (Walker et al., 2017). We further showed that administration of GDF11 *in vivo* more potently induces SMAD phosphorylation in the myocardium compared to GDF8 (Walker et al., 2017). These differences implicate residue differences between GDF11 and GDF8, particularly those clustered around the type I binding interface, in determining signaling potency, likely via effects on dimer stability and stability of receptor interactions. Consistent with this possibility, structural analysis and mutational studies of the ternary complex of GDF11 with type I receptor Alk5 and type II receptor ActRIIB revealed that different mechanisms regulate specificity and binding with type I receptor, compared to TGFβ, providing an explanation for how GDF11 and the TGFβ activin class more effectively facilitate low-affinity type I interactions (Goebel et al., 2019). These biochemical and structural studies indicate that GDF11 and GDF8 are unlikely to be functionally equivalent, especially when ligand concentrations are low, as they typically are *in vivo* (Walker et al., 2017; Goebel et al., 2019).

In this study, we evaluate GDF11 and GDF8 functional equivalence *in vivo* by utilizing the CRISPR/Cas9 system to introduce GDF8-like amino acid substitutions into GDF11 and GDF11-like substitutions into GDF8. These sequence alterations in GDF11, which previously were shown to diminish potency of the resulting protein (Walker et al., 2017), caused a perturbation of the axial skeletal structure of mutant mice during development that persists into adulthood. In contrast, the sequence alterations introduced into GDF8, which previously were shown to increase the potency of the resulting ligand (Walker et al., 2017), did not produce observable developmental phenotypes. As such, we generated a third line of mutant animals, in which the entire GDF8 mature domain was replaced with the corresponding mature domain sequence of GDF11, resulting in full replacement of the endogenous GDF8 signaling domain with that of GDF11. These mature domain (MD) mutants had up to 50-fold greater levels of GDF11 in circulation, with concomitant depletion of GDF8 to undetectable levels and showed modestly decreased skeletal muscle mass, with no apparent impact on postnatal survival, total adult body weight, or the development and function of other organ systems.

While we were performing our study, the Se-jin Lee group published a study that used a similar genetic approach as our third mouse line to replace the *Mstn* gene sequences encoding the mature C-terminal peptide with the full mature domain of *Gdf11* (Lee et al., 2022). Their characterization of these mice (Lee et al., 2022) supports our data shown here, showing that GDF11 entirely replaced circulating MSTN and increased GDF11 levels ∼30-40-fold (Lee et al., 2022). However, our findings extend these observations, addressing the converse hypothesis as well and showing that diminution of GDF11 potency—through targeted replacement of two key amino acids from GDF8—causes a significant developmental defect in osteogenesis, distinct from that seen with modulation of GDF8. Taken together, our findings elucidate precise, differential molecular mechanisms underpinning the biological actions of GDF11 and GDF8 that cannot be explained solely by differences in *in vivo* ligand concentrations and patterns of expression. They also provide direct evidence that structural and biochemical differences in these ligand’s mature signaling domains contribute significantly to their unique roles in mammalian development and organ physiology.

## Results

### Generation and characterization of chimeric amino acid GDF11 and GDF8 mice

We previously reported that substitution of two residues located in the fingertip of GDF11 (D89 and Q91) into the analogous region of GDF8 (in place of G89 and E91) enhanced SMAD signaling activity of the hybrid GDF8 molecule by approximately 50% (Walker et al., 2017). This result indicates that sequence differences in the mature GDF11 and GDF8 proteins are likely responsible for differences in ligand signaling and function. To address whether such sequence-determined signaling differences impact *in vivo* activities of GDF11 and GDF8, we used CRISPR/Cas9 to create two lines of chimeric mice **(Strategy 1, Fig. 1C**), in which we replaced D89 and Q91 residues within the *Gdf11* locus with the analogous G89 and E91 residues from *Gdf8*, or conversely, replaced G89 and E91 within native *Gdf8* with D89 and Q91 from *Gdf11*. The third chimeric line we generated replaced the full mature domain region within the *Gdf8* locus with the corresponding region from *Gdf8* **(Strategy 2, Fig. 1D**). Based on our prior *in vitro* studies, substitution of all the unique GDF11 residues into GDF8 in this manner is able to enhance signaling of the resulting protein to be approximately 5-fold more potent than wild-type GDF8 (Walker et al., 2017).

To generate the two amino acid modified lines, we constructed genetically modified *Gdf11* **(Fig. S1A)** and *Gdf8* **(Fig. S1B)** single-stranded DNA (ssDNA) donor plasmids containing the mutant codons, flanked by ∼80bp homologous arms. In generating the full mature domain replacement line, we constructed a genetically modified *Gdf8* double-stranded DNA (dsDNA) donor plasmid **(Fig. S1C)** containing the GDF11 mature domain sequences, flanked by ∼4kb homologous arms. After homologous recombination in embryonic stem cells, targeted microinjections into C57BL/6J zygotes, and implantation of zygotes into C57BL/6J surrogate females, we produced F0 founders with the chimeric allele incorporated in the germline. The ssDNA donor template incorporating *Gdf8*-like G89 and E91 residues into the native *Gdf11* locus also introduced an *AseI* restriction enzyme unique to *Gdf8* **(Fig. S1A)** as a genetic marker for downstream genotyping. Likewise, the ssDNA donor template incorporating *Gdf11*-like D89 and Q91 residues into the native *Gdf8* locus **(Fig. S1B)** and the dsDNA donor template containing the *Gdf11* full mature domain sequences **(Fig. S1C)** removed the same *AseI* site from the *Gdf8* locus. We verified successful integration of the chimeric constructs at the *Gdf11* and *Gdf8* loci, respectively, via Sanger sequencing (**Fig. S1A, B, C**) and by PCR validation and subcloning (**Fig. S1G, H, I**), which also confirmed the presence of the unique *AseI* restriction enzyme site in the modified *Gdf11* locus and absence of the same *AseI* site at the modified *Gdf8* loci. We further confirmed integration of silent mutations included in the donor templates, whose purpose was to mutate the PAM sequence to prevent further cutting after donor construct integration (**Fig. S1A, B, C**). Collectively, these results validated our genetic modification strategy to integrate *Gdf11*-like and *Gdf8*-like changes into the *Gdf8* and *Gdf11* loci, respectively.

Through this process, we successfully generated:

1. *Gdf11^Gdf8aa^* mice (with *Gdf8* amino acid residues G89 and E91 replacing the corresponding residues in *Gdf11*) **(Fig. S1A)** a. Mono-allelic (*Gdf11^+/8aa^*) / bi-allelic (*Gdf11^8aa/8aa^*) / wild-type (*Gdf11^+/+^*)
2. *Gdf8^Gdf11aa^* mice (with *Gdf11* amino acid residues D89 and Q91 replacing the corresponding residues in *Gdf8*) **(Fig. S1B)** a. Mono-allelic (*Gdf8^+/11aa^*) / bi-allelic (*Gdf8^11aa/11aa^*) / wild-type (*Gdf8^+/+^*)
3. *Gdf8^Gdf11MD^* mice (with the *Gdf11* mature domain replacing the *Gdf8* mature domain in the *Gdf8* locus) (Fig. S1C) a. Mono-allelic (*Gdf8^+/11MD^*) / bi-allelic (*Gdf8^11MD/11MD^*) / wild-type (*Gdf8^+/+^*)

We backcrossed the knock-in alleles five generations (F5) in each chimeric line prior to analyses to breed out potential off-target modifications and confirmed Mendelian ratios of allele inheritance to rule out potential embryonic lethality resulting from the genetic modifications (**Table S1**). Targeted Locus Amplification (TLA) sequencing (de Vree et al., 2014), performed on bone marrow DNA harvested from F5 mono-allelic and bi-allelic offspring from *Gdf11^Gdf8aa^* **(Fig. S2A, B)** *Gdf8^Gdf11aa^* **(Fig. S2C, D)**, and *Gdf8^Gdf11MD^* **(Fig. S2E, F)** mice and aligned to the mouse mm10 genome sequence, further confirmed correct integration of the desired mutant sequences into the native *Gdf11* and *Gdf8* loci, with no evidence across the whole genome of structural variation surrounding the integration site or within the insert, of incorrect or off-target integration events, or of a locus duplication at the integration site matching the wild-type allele. Taken together, these results indicate that the *Gdf11^Gdf8aa^* and *Gdf8^Gdf11aa^* chimeric lines successfully integrated the intended *Gdf8* or *Gdf11* nucleotide alterations leading to the anticipated amino acid changes, with no other genomic off-target mutations detected, and that the *Gdf8^Gdf11MD^* line successfully integrated the full mature domain sequences of *Gdf11* and replaced the native mature domain region of *Gdf8*, with no evidence of incorrect targeting.

### Circulating GDF11 concentration increases 50-fold in *Gdf8^Gdf11MD^* mutants, while GDF11 and GDF8 levels in *Gdf11^Gdf8aa^* and *Gdf8^Gdf11aa^* mutants remain unchanged

To determine whether full replacement of the mature domain of native *Gdf8* with *Gdf11* altered circulating protein levels *in vivo*, we collected serum from *Gdf8^+/+^*, *Gdf8^+/11MD^*, and *Gdf8^11MD/11MD^* mice at 10-14 weeks of age and measured endogenous GDF11 and GDF8 levels, using a liquid chromatography tandem mass spectrometry assay that distinguishes between GDF11 and GDF8 by detecting two differential peptide fragments between the two proteins (Garbern et al., 2019) (**Fig. 2**). In the bi-allelic *Gdf8^11MD/11MD^* mutants (n=10), circulating GDF11 concentrations increased approximately 50-fold above normal levels (**Fig. 2A**), while circulating GDF8 concentrations decreased below level of detection (<LOD) (**Fig. 2B**). The same trend in GDF11 and GDF8 concentrations was observed in male and female mice separately, with significant differences in ligand concentration varying according to allelic dosage across the three genotypes (*Gdf8^+/+^*, n=10; *Gdf8^+/11MD^*, n=8; and *Gdf8^11MD/11MD^*, n=10). The levels of GDF11 peptides measured in *Gdf8^11MD/11MD^* mutants (**Fig. 2A**) was comparable to the levels of GDF8 in *Gdf8^+/+^* mice (**Fig. 2B**) and to the combined concentrations of GDF11 + GDF8 in *Gdf8^+/11MD^* mice (**Fig. 2A,B**), suggesting a direct replacement of mature GDF11 for GDF8, with expression levels ultimately determined by either cis-regulatory elements in the *Gdf8* locus or association with the GDF8 prodomain, or a combination of these two factors. In support of this interpretation, the total pool of GDF11 + GDF8 did not differ for any of the chimeric genotypes (**Fig. 2C**). Mass spectrometry data from the Se-jin Lee group of the *Gdf8* full coding region-replaced chimeras corroborate our findings (Lee et al., 2022). They also reported no detectable MSTN and a ∼30-40-fold increase in circulating GDF11 in *Mstn^Gdf11/Gdf11^* mice (Lee et al., 2022), with *Mstn^+/Gdf11^* mice having intermediate levels of the two ligands (Lee et al., 2022). Our study further expands this analysis of the impact of mature domain sequence on ligand expression by measuring endogenous GDF11 and GDF8 in serum from *Gdf11^Gdf8aa^* and *Gdf8^Gdf11aa^* mutants as well, showing that serum GDF11 and GDF8 protein concentrations are not significantly altered in either mono-allelic or bi-allelic *Gdf11^Gdf8aa^* or *Gdf8^Gdf11aa^* mutants, compared to wild-type mice (**Fig. 2A, B, C**). These data indicate that the dual amino acid substitutions alone did not impact endogenous expression or circulation of GDF11 or GDF8. Thus, despite some reports that high levels of GDF11 in humans are associated with adverse health consequences (Egerman et al., 2015), our data, together with those of Lee and colleagues (Lee et al., 2022), indicate that GDF11 can rise to extremely high levels *in vivo* without apparent negative health consequences or premature death.

**Figure 2.**
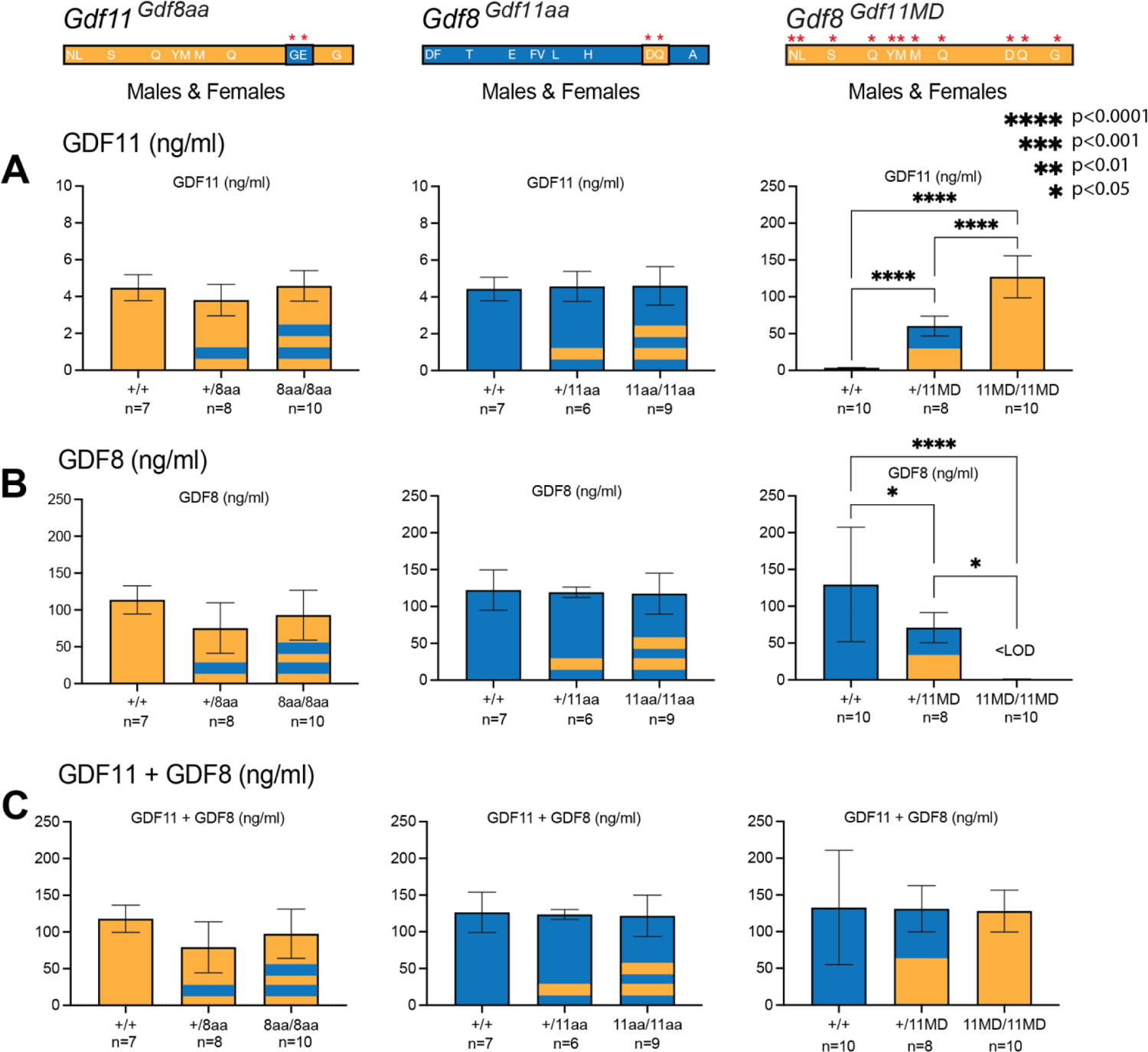
GDF11 and GDF8 serum concentrations in *Gdf11^Gdf8aa^*, *Gdf8^Gdf11aa^*, and *Gdf8^Gdf11MD^* mice. Liquid chromatography tandem mass spectrometry measurements of GDF11 (**A**) and GDF8 (**B**) in *Gdf11^+/+^* (n=7), *Gdf11^+/8aa^* (n=8), and *Gdf11^8aa/8aa^* (n=10) mice **(left plots),** in *Gdf8^+/+^* (n=7), *Gdf8^+/11aa^* (n=6), and *Gdf8^11aa/11aa^* (n=9) mice **(middle plots)**, and in *Gdf8^+/+^* (n=10), *Gdf8^+/11MD^* (n=8), and *Gdf8^11MD/11MD^* (n=10) mice. <LOD, below the level of detection. The same trends were observed in male and female mice separately. (**C**) Combined concentration of GDF11 and GDF8 in *Gdf11^Gdf8aa^*, *Gdf8^Gdf11aa^*, and *Gdf8^Gdf11MD^* mice as a measure of total ligand levels. Statistical analyses performed by one way ANOVA with Tukey’s correction for multiple comparisons. Amino acid residues of *Gdf11* represented in orange and *Gdf8* in blue. For *Gdf11^Gdf8aa^, Gdf8^Gdf11aa^* lines, 1 stripe denotes mono-allelic replacement, 2 stripes, bi-allelic replacement. For *Gdf8^Gdf11MD^* mice, half orange denotes mono-allelic replacement, full orange denotes bi-allelic replacement.

### GDF11 dampening in bi-allelic *Gdf11^Gdf8aa^* mutant embryos recapitulates developmental phenotype seen with *Gdf11* loss-of-function

Germline deletion of *Gdf11* results in perinatal lethality, and both homozygous (*Gdf11^-/-^*) and heterozygous (*Gdf11^+/-^*) disruption of *Gdf11* causes developmental abnormalities of the skeleton (McPherron et al., 1999; Walker et al., 2016)—notably the formation of extra thoracic vertebrae—and kidney agenesis in pups (Esquela & Lee, 2003; McPherron et al., 2009). We therefore assessed early-stage skeletal development in our chimeric embryos (**Fig. 3**). Six sets of F4 mono-allelic mutant males were bred with mono-allelic mutant females within each *Gdf11^Gdf8aa^*, *Gdf8^Gdf11aa^*, and *Gdf8^Gdf11MD^* mouse line to generate F5 embryos that were harvested on embryonic day 18.5 (E18.5), eviscerated, and stained with alizarin red and alcian blue (**Fig. 3**). Prior to evisceration, tissue samples from the posterior skin of prenatal pups were collected for genotyping by PCR validation and sub-cloning. Spleen and liver samples were also taken from each embryo for genotyping (data not shown) to confirm that maternal DNA from the fallopian tubes and gestational sacs would not obfuscate genotyping results by contaminating the collected embryonic tissue.

**Figure 3.**
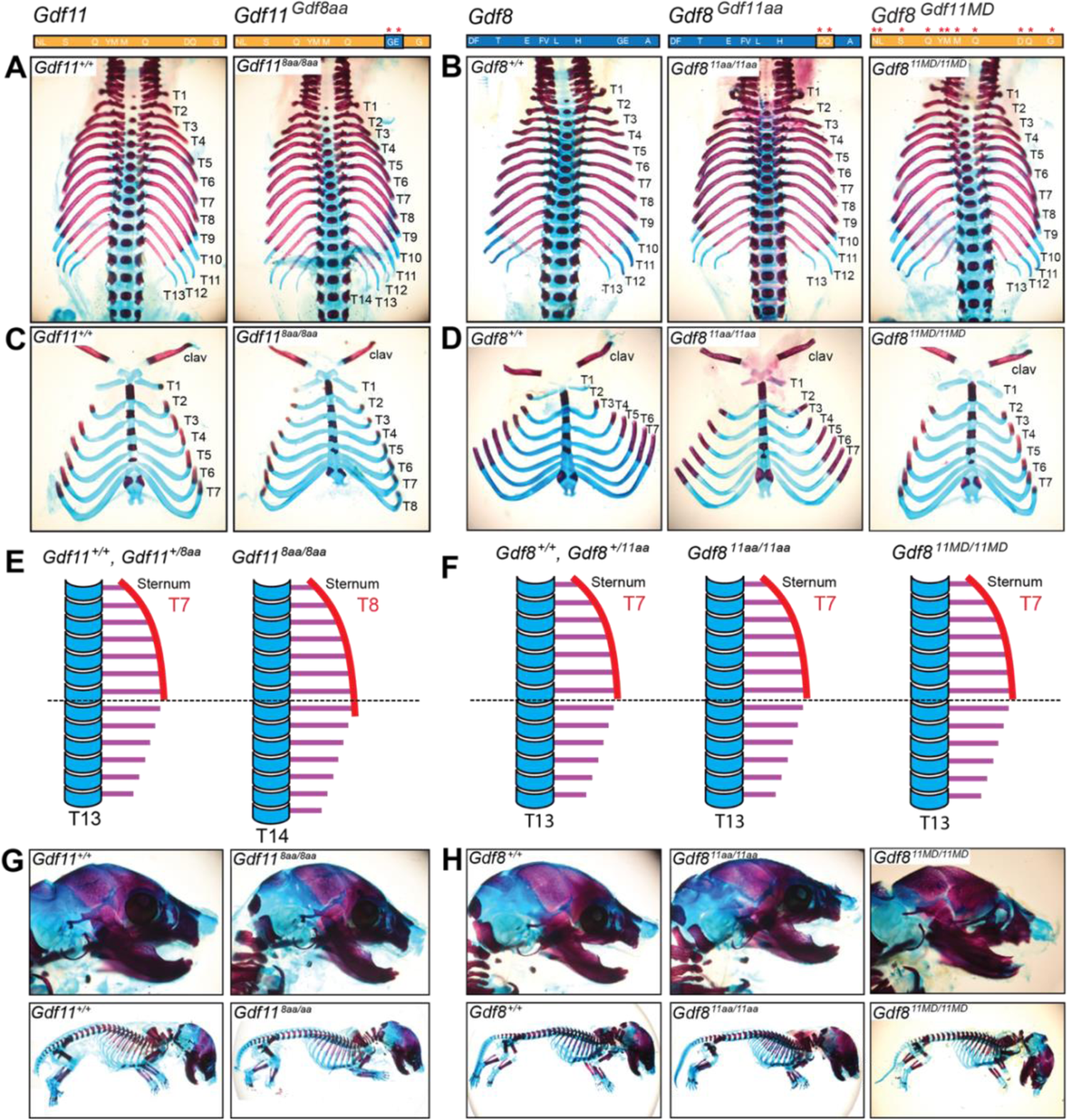
Axial skeletal patterning, craniofacial bone, and limb development in *Gdf11^Gdf8aa^*, *Gdf8^Gdf11aa^*, and *Gdf8^Gdf11MD^* mice. **(A, B)** Skeletons from *Gdf11^Gdf8aa^*, *Gdf8^Gdf11aa^*, and *Gdf8^Gdf11MD^* embryos harvested at E18.5 were stained with alizarin red and alcian blue. *Gdf11^+/+^* (leftmost), *Gdf11^8aa/8aa^* (2nd from left), *Gdf8^+/+^* (center), *Gdf8^+/11aa^* (2nd from right), and *Gdf8^11MD/11MD^* (rightmost) vertebral columns, vertebrosternal ribs, cranofacial bones, and full skeletons are shown. *Gdf11^8aa/8aa^* skeletons exhibited abnormal axial vertebral patterning, having T14 total axial vertebrae, whereas all other mutants presented with T13 total axial vertebrae. **(C)** *Gdf11^+/+^* skeletons also exhibited T8 vertebrosternal ribs, compared to T7 vertebrosternal ribs in all other mutants **(D)**. **(E)** Schematic of axial vertebral columns observed in *Gdf11^Gdf8aa^* embryos. *Gdf11^+/8aa^* mutants also had T13 total axial vertebrae and T7 vertebrosternal ribs. **(F)** Schematic of axial vertebral columns observed in *Gdf8^Gdf11aa^* and *Gdf8^Gdf11MD^* embryos. *Gdf8^+/11aa^* and *Gdf8^+/11MD^* mutants had T13 total axial vertebrae and T7 vertebrosternal ribs. **(G)** The craniofacial bone and overall skeletal shape of *Gdf11^8aa/8aa^*, *Gdf8^11aa/11aa^*, *Gdf8^11MD/11MD^* mutants were indistinguishable from *Gdf11^+/+^* and *Gdf8^+/+^* mice, respectively. The limbs and digits of the mutant mice from both lines were also similar to wild-type. Schematic of the amino acid residues in *Gdf11* (orange) and *Gdf8* (blue) locus.

Upon imaging the skeletons, we discovered distinct skeletal transformations in the *Gdf11^Gdf8aa^* mouse line. Specifically, the bi-allelic *Gdf11^8aa/8aa^* mutants exhibited one extra thoracic vertebra, with T14 axial vertebrae in total, compared to T13 vertebrae in wild-type mice (**Fig. 3A, E**). We also observed an extra vertebrosternal rib in *Gdf11^8aa/8aa^* mutants, resulting in a total of T8 ribs connected to the sternum, compared to T7 ribs in *Gdf11^+/+^* and *Gdf11^+/8aa^* mice (**Fig. 3C, E)**. Phenotypically, this differential development directly compares to the axial skeletal phenotype observed in heterozygous *Gdf11^+/-^* mice, which also have 14 total thoracic vertebrae and 8 pairs of ribs fused to the sternum (McPherron et al., 1999). In contrast, homozygous *Gdf11^-/-^* knockout mice have 17-18 total thoracic vertebrae and 10-11 pairs of vertebrosternal ribs (McPherron et al., 1999). We further verified that the malformation documented occurred only in *Gdf11^8aa/8aa^* mutants, and with 100% penetrance (**Table 1**). Moreover, given that circulating ligand levels in these chimeric mice remained unaltered, the observed differential skeletal phenotypes are specifically attributable to the dual amino acid changes made in the protein sequences and structures—not to alterations of circulating protein levels. In contrast, the *Gdf8^Gdf11aa^* mutant skeletons appeared indistinguishable from wild-type (**Fig. 3H**), and none of the mono-allelic or bi-allelic mutants produced a measurable skeletal phenotype (**Fig. 3B, D, F, Table 1**). As stated previously, we generated the *Gdf8^Gdf11MD^* mutant line to investigate whether increasing GDF8 potency to the maximum level of GDF11, by replacing the entire mature domain of GDF8 with that of GDF11, would result in observable phenotypic outcomes. However, axial skeletal analysis of *Gdf8^Gdf11MD^* mice also revealed no measurable defects in the mutant skeletons, compared to wild-type (**Fig. 3B, D, F, Table 1**). Reported analyses of the related *Mstn^Gdf11/Gdf11^* line generated by Lee and colleagues similarly found no abnormalities of axial skeletal patterning, though some decrement in bone density and alterations in trabeculae were noted, almost exclusively in males (Lee et al., 2022). These authors did not evaluate the impact of the dual amino acid substitutions reported here.

**Table 1.**
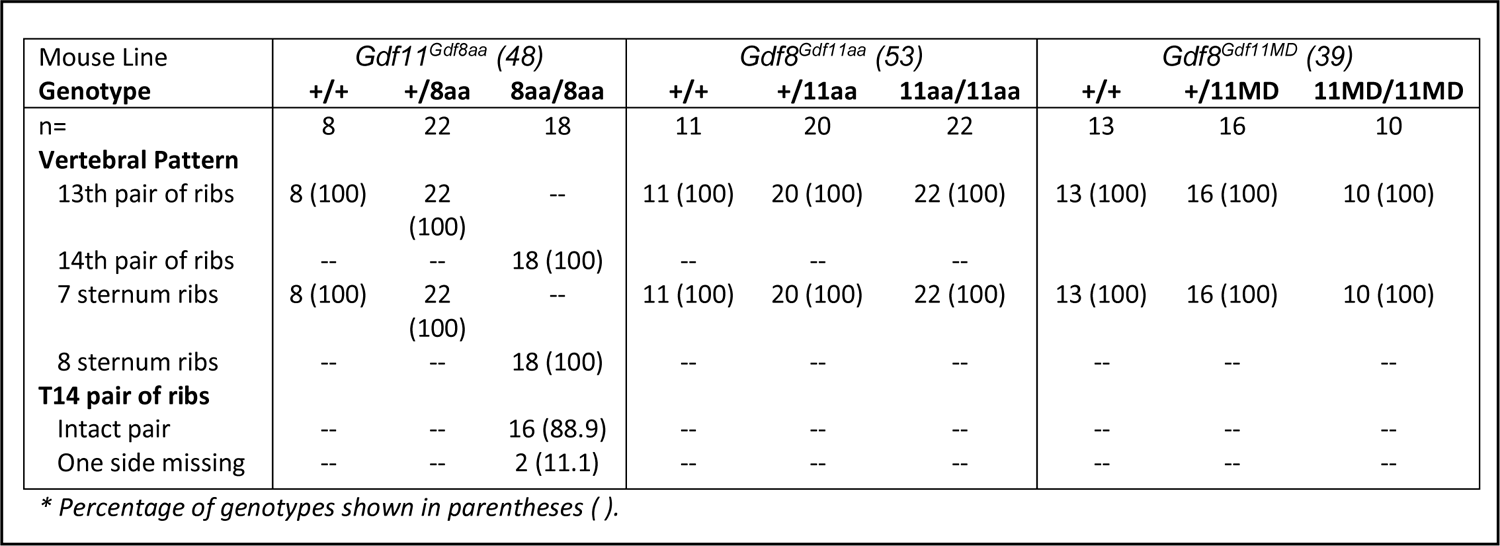
Skeletal analysis *Gdf11^Gdf8aa^, Gdf8^Gdf11aa^*, and *Gdf8^Gdf11MD^* mouse embryos in C57BL/6J background. Comparison of vertebral columns and vertebrosternal ribs from embryos harvested at E18.5. In *Gdf11^Gdf8aa^* mice, T14 total axial vertebrae and T8 vertebrosternal ribs were observed in *Gdf11^8aa/8aa^* embryos with 100% penetrance, compared to T13 vertebrae and T7 vertebrosternal ribs in 100% of *Gdf11^+/+^* and *Gdf11^+/8aa^* embryos. *Gdf8^Gdf11aa^* and *Gdf8^Gdf11MD^* embryos all had T13 axial vertebrae and T7 vertebrosternal ribs with 100% penetrance. No additional anomalies outside of these vertebral aberrations were observed. The total number of embryos represents two separate cohorts harvested from multiple F4 mono-allelic *Gdf11^Gdf8aa^, Gdf8^Gdf11aa^*, and *Gdf8^Gdf11MD^* breeding pairs.

We also investigated early-stage craniofacial bone development in *Gdf11* and *Gdf8* mutant mice (**Fig. 3G, H)**, since palatal defects have been reported in mice with *Gdf11^-/-^* deletion (McPherron et al., 1999; Cox et al., 2019) and in humans with *Gdf11* loss-of-function alleles (Ravenscroft et al., 2021). In our inspection, the only skeletal differences were detected in the axial vertebral patterning and vertebrosternal rib count of *Gdf11^8aa/8aa^* mutant mice (**Fig. 3A, C)**. No defects were noted in the limbs or cranium of *Gdf11^Gdf8aa^, Gdf8^Gdf11aa^*, or *Gdf8^Gdf11MD^* mutants, compared to *Gdf11^+/+^* and *Gdf8^+/+^* mice (**Fig. 3G, H)**. We also saw no palatal defects consistent with those previously reported for Gdf11-null mice (McPherron et al., 1999), nor did we observe a hole in the otic capsule of mutant mice, which has been reported in *Gdf11* indel and gene-targeted mice (Goldstein et al., 2019) (**Fig. 3G, H)**. These results suggest that while full potency of GDF11 may not be necessary for craniofacial bone development, it is crucial for proper axial skeletal development. Therefore, dampening the potency of mature GDF11 with substitution of GDF8 residues is not compatible with maintaining fully normal developmental function, even with appropriate patterning of expression provided by the endogenous *Gdf11* genomic locus. On the other hand, it appears that increasing the potency of GDF8, even to the maximum level of GDF11, does not elicit malformations detectable in early development. Together, these data underscore the notion that GDF11 and GDF8 are functionally distinct during development.

### *Gdf8^Gdf11MD^* mutants exhibit decreased skeletal muscle mass, while the muscles of *Gdf11^Gdf8aa^* and *Gdf8^Gdf11aa^* mutants are not significantly altered

Next in our characterization, we examined early-stage skeletal muscle and cardiac development in the *Gdf11^Gdf8aa^, Gdf8^Gdf11aa^*, and *Gdf8^Gdf11MD^* mouse lines. Prior studies indicate that genetic inactivation of *Gdf8* dramatically increases muscle mass and alters fiber type distribution across multiple animal species and in a dose-dependent manner (McPherron et al., 1997; McPherron & Lee, 1997), whereas boosting levels of GDF8 protein has been shown to drive muscle wasting (Zimmer et al., 2002; Stolz et al., 2008). We therefore sought to determine whether enhancing the potency of mature GDF8 in the *Gdf8^Gdf11aa^* or *Gdf8^Gdf11MD^* mutants might reduce muscle mass compared to *Gdf8^+/+^* mice, harvesting and analyzing the wet weight of the tibialis anterior (TA) (**Fig. 4C, D)**, quadriceps (**Fig. 4F, G)**, and triceps (**Fig. 4I, J)** muscles postmortem across all three chimeric lines at 10-14 weeks of age. Muscle weights were normalized to both total body weight (**Fig. 4A, B)** and to tibia bone length (**Fig. S3**) in the *Gdf11^Gdf8aa^, Gdf8^Gdf11aa^*, and *Gdf8^Gdf11MD^* mutants (**Fig. 4E, H, K**) to assess possible changes in muscle mass. Additionally, we harvested and weighed the kidneys from each mouse and normalized them to total body weight (**Fig. S4**).

**Figure 4.**
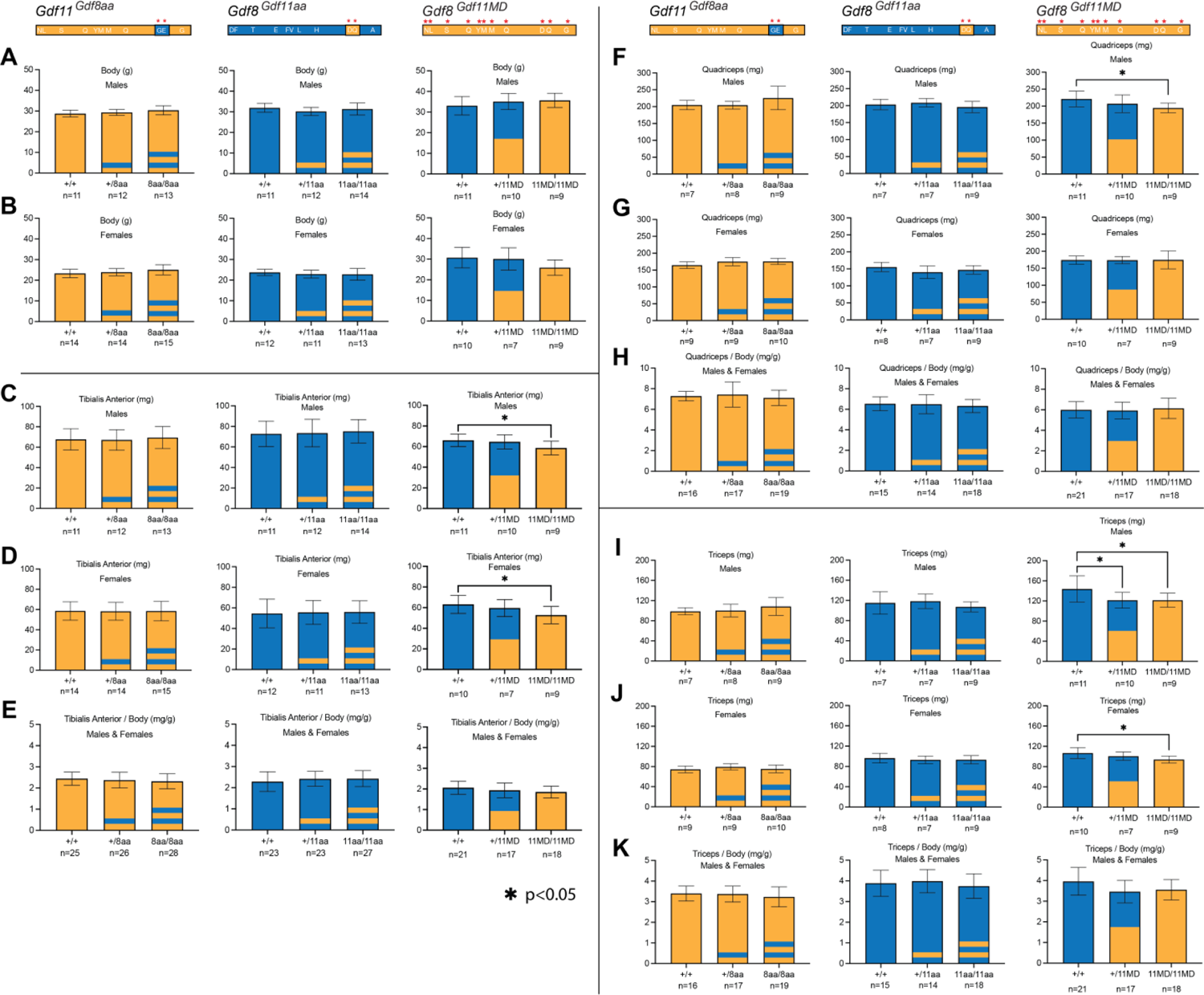
Body weight and muscle mass in *Gdf11^Gdf8aa^, Gdf8^Gdf11aa^*, and *Gdf8^Gdf11MD^* mice. Tibialis anterior (TA) **(C, D)**, quadriceps **(F, G)**, and triceps **(I, J)** muscles were collected postmortem from *Gdf11^Gdf8aa^, Gdf8^Gdf11aa^*, and *Gdf8^Gdf11MD^* mice at 10-14 weeks of age. Mice were weighed prior to dissection **(A, B)**, and weights of the TA, quadriceps, and triceps muscles were normalized to body weight **(E, H, K)**. Statistically significant decreases in TA **(C, D)**, quadriceps **(F)**, and triceps **(I, J)** muscle mass were recorded in male and female *Gdf8^11MD/11MD^* mice, compared to *Gdf8^+/+^*. However, no significant differences were observed for normalized muscle weights in these mutant mice, compared to wild-type. Statistical analysis was performed by one way ANOVA with Tukey’s correction for multiple comparisons. The amino acid residues of *Gdf11* are represented in orange and *Gdf8* in blue. For *Gdf11^Gdf8aa^* and *Gdf8^Gdf11aa^* lines, 1 stripe and 2 stripes denote mono-allelic and bi-allelic replacement, respectively. For *Gdf8^Gdf11MD^* mice, half orange denotes mono-allelic replacement, while full orange denotes bi-allelic replacement. Also see Figures S3 and S4.

Across the chimeric amino acid *Gdf8^Gdf11aa^* and *Gdf11^Gdf8aa^* lines, we saw no significant differences in body weight (**Fig. 4A, B)**, TA (**Fig. 4C, D)**, quadriceps (**Fig. 4F, G)**, or triceps weights (**Fig. 4I, J)**. Skeletal muscle weights normalized to total body weight (**Fig. 4E, H, K**) and to tibia length **(Fig S3B, C, D)** were indistinguishable. By contrast, the weights of these skeletal muscles were significantly decreased across the *Gdf8^Gdf11MD^* mutants (**Fig. 4C, D, F, I, J)**. Notably, TA (**Fig. 4C**), quadriceps (**Fig. 4F**), and triceps (**Fig. 4I**) weights all were reduced to a statistically significant level in bi-allelic *Gdf8^11MD/11MD^* mutant males (n=9, p<0.05), compared to sex-matched *Gdf8^+/+^* mice (n=10), with similarly significant decreases recorded in TA (**Fig. 4D**) and triceps (**Fig. 4J**) weights in *Gdf8^11MD/11MD^* mutant females (n=9, p<0.05) as well. While all three muscle groups in *Gdf8^11MD/11MD^* mice normalized to overall body weight did not yield statistically significant differences (**Fig. 4E, H, K**), the raw weights of TA muscle normalized to tibia length (n=18, p<0.01) **(Fig. S3B)** and triceps muscle normalized to tibia length (n=18, p<0.05) **(Fig. S3D)** did show statistically significant decreases, compared to *Gdf8^+/+^* mice (n=21). In all cases, separation by males and females resulted in shifts in the mean muscle mass between the sexes. Analysis of muscle mass in *Mstn^Gdf11/Gdf11^* mice showed similar (∼10%) reductions in muscle mass in male, but not female, mice in which the mature domain of GDF8 was replaced by that of GDF11, with no differences in fiber composition (Lee et al., 2022). However, these studies did not include analysis of the dual amino acid substitutions reported here.

While GDF11 loss-of-function frequently leads to kidney agenesis (Esquela & Lee, 2003; McPherron et al., 2009), we did not observe any chimeric mouse lacking a kidney. The combined weight of both kidneys (**Fig. S4A, B, C**) also did not show any significant change in *Gdf11^Gdf8aa^, Gdf8^Gdf11aa^*, and *Gdf8^Gdf11MD^* mutants, compared to wild-type mice. Interestingly, we did record a statistically significant decrease in the liver weight of *Gdf8^11MD/11MD^* females (n=9), compared to *Gdf8^+/+^* females (n=10, p<0.01) **(Fig. S4E)**, but a similarly significant decrease was not found in *Gdf8^Gdf11MD^* mutant males **(Fig. S4D)**, despite a modest, yet progressive, weight decline in the liver of *Gdf8^+/11MD^* mutants, followed by *Gdf8^11MD/11MD^* mutants, compared *Gdf8^+/+^* mice (**Fig. S4D, E, F**). This trend was not seen in *Gdf8^Gdf11aa^* mice (**Fig. S4D, E, F**).

Taken together, these data indicate that the increased potency conferred to GDF8 by substitution of the GDF11 mature domain significantly impacted skeletal muscle mass in young *Gdf8^Gdf11MD^* mutant mice, in line with prior reports that endogenous GDF8 negatively regulates muscle development. The observed effects were distinct from those conferred to young *Gdf8^Gdf11aa^* mutant mice by substitution of only the *Gdf11*-like 89/91 residues—which did not yield any significant change in skeletal muscle mass. These data indicate a clear difference in potency between the mature domains of GDF11 and GDF8, as well as between the full GDF11 mature domain-replaced ligand and the double amino acid-substituted ligand generated here. Additionally, though we did not observe signs of kidney agenesis in the *Gdf8^Gdf11MD^* mouse line, we recorded a trending decline in liver weight, most significant in *Gdf8^Gdf11MD^* mutant females. These data suggest a possible systemic effect on the liver—which does not produce GDF8—that could reflect a direct effect on liver hepatocytes or, more likely, be connected to the local GDF8 regulation of skeletal muscle mass. In such a scenario, skeletal muscle may be more sensitive to increased potency of GDF8, and as a result, a change in muscle mass indirectly affects liver development or homeostasis.

### Baseline cardiac physiology and function remains unchanged in *Gdf11^Gdf8aa^, Gdf8^Gdf11aa^*, and *Gdf8^Gdf11MD^* mutants

Previous studies have shown that administration of exogenous recombinant GDF11 to aged mice reduces cardiac hypertrophy (Loffredo et al., 2013), and fetal cardiac GDF8 has also been implicated in early-stage heart development (Sharma et al., 1999). Therefore, we harvested and weighed the hearts of all chimeric mice at 10-14 weeks of age (**Fig. 5A, B)**. In the *Gdf11^Gdf8aa^* line, we found a statistically significant increase in heart weight of *Gdf11^8aa/8aa^* males (n=13), compared to *Gdf11^+/+^* males (n=11, p<0.05) and to *Gdf11^+/8aa^* males (n=12, p<0.05) (**Fig. 5A**). However, this difference was not observed in *Gdf11^Gdf8aa^* female mice (**Fig. 5B**), and the overall weight of *Gdf11^Gdf8aa^* mutant hearts normalized to body weight did not show a significant difference (**Fig. 5C**). Analyses of *Gdf8^Gdf11aa^* and *Gdf8^Gdf11MD^* mouse lines showed no significant difference in the weight of mutant hearts, compared to that of *Gdf8^+/+^* mice (**Fig. 5A, B)**. Further histopathological analysis of cardiac tissue also did not reveal significant differences in cardiomyocyte cross-sectional area (CSA) in any mutant lines compared to age- and sex-matched *Gdf11^+/+^* and *Gdf8^+/+^* mice (**Fig. 5D**). We also investigated whether the mutations introduced to native *Gdf11* and *Gdf8* altered baseline cardiac physiology or function by performing blinded echocardiographic studies on all the chimeric mice at 10-14 weeks of age (**Fig. 5E, F)**. In all three lines, echocardiographic imaging indicated equivalent baseline cardiac function. Fractional shortening (FS) (**Fig. 5E**) and left ventricular heart dimensions (LVAW; LVPW; LVID) during systole and diastole (**Fig. 5F**) were consistent across all genotypes, with no significant differences observed. Cardiac ejection fraction (ES) was also comparable across all three lines (**Fig. S5A, B, C**).

**Figure 5.**
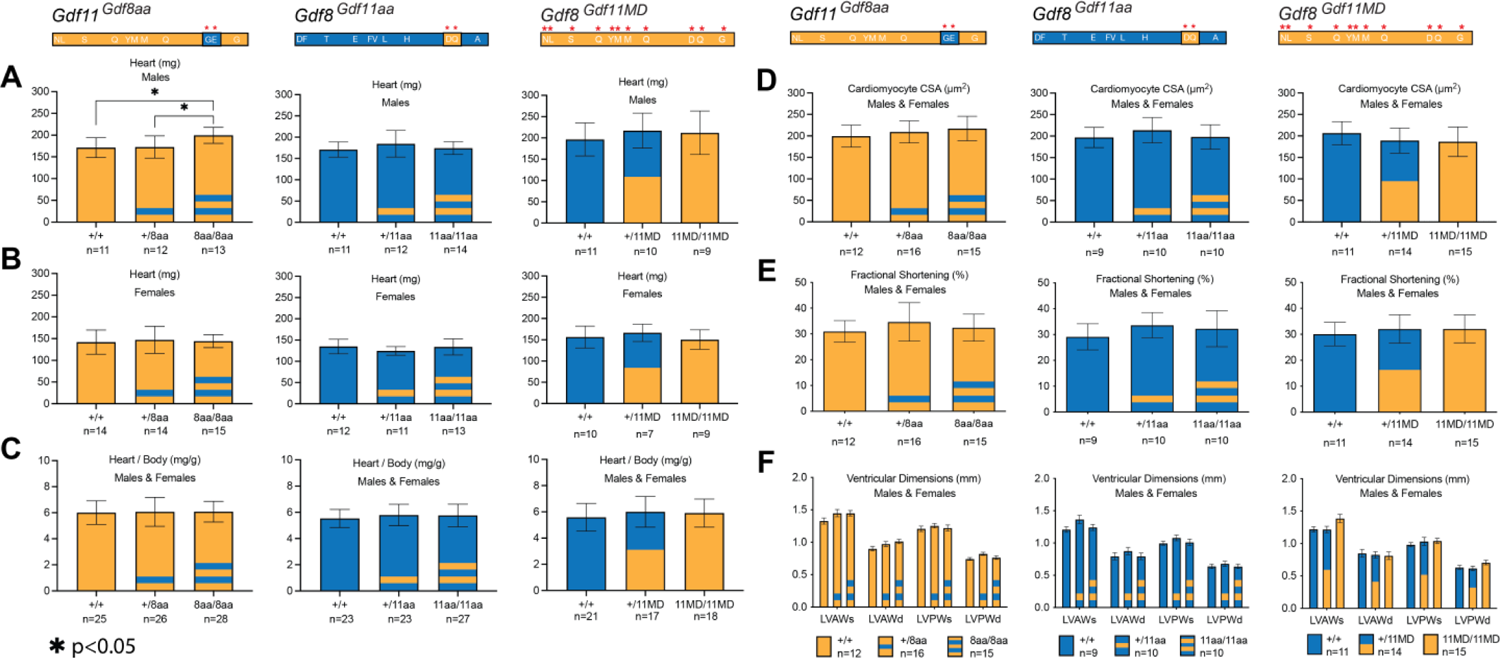
Baseline heart physiological and functional measurements in *Gdf11^Gdf8aa^, Gdf8^Gdf11aa^*, and *Gdf8^Gdf11MD^* mice. The hearts of *Gdf11^Gdf8aa^, Gdf8^Gdf11aa^*, and *Gdf8^Gdf11MD^* male and female mice were harvested and weighed at 10-14 weeks of age **(A, B)** and normalized to overall body weight **(C)**. Cardiomyocyte cross-sectional area (CSA) **(D)**, fractional shortening (FS) **(E)**, and left ventricular heart dimensions (LVAW; LVPW; LVID) **(F)** during systole and diastole were also measured across all three mouse lines. A statistically significant increase in heart weight was found in *Gdf11^8aa/8aa^* males (n=13) **(A)**, compared to *Gdf11^+/+^* (n=11, p<0.05) and to *Gdf11^+/8aa^* males (n=12, p<0.05). However, no similar difference was observed in *Gdf11^Gdf8aa^* females **(B)**, and the overall weight of *Gdf11^Gdf8aa^* mutant hearts normalized to body weight **(C)** did not yield a significant difference either. The cross-sectional area of cardiomyocytes was measured using FIJI software (scale bar=100μm). M-mode was used to measure left ventricular interventricular septal wall thickness (IVS/LVAW), left ventricular posterior wall thickness (LVPW), and left ventricular internal diameter (LVID). Statistical analysis was performed by one way ANOVA with Tukey’s correction for multiple comparisons. Amino acid residues of *Gdf11* represented in orange and *Gdf8* in blue. Also see Figure S5.

These results indicate that the *Gdf11^Gdf8aa^, Gdf8^Gdf11aa^*, and *Gdf8^Gdf11MD^* mutant hearts have function and physiology similar to wild-type mice. While no heart weight phenotype has been reported for *Gdf11^+/-^* mice, a difference was found in the heart weight of bi-allelic *Gdf11^8aa/8aa^* mutant males (**Fig. 5A**). However, this difference was not present in *Gdf11^8aa/8aa^* female hearts (**Fig. 5B**), and overall, we saw no significant differences across heart physiology and function of *Gdf8^Gdf11aa^* mutants. Furthermore, conferring the full potency of GDF11 to GDF8 did not significantly impact cardiac muscle cross sectional area or ventricular dimensions in young *Gdf8^Gdf11MD^* mutant mice (**Figs. 5, S5)**. If potency changes in either GDF11 or GDF8 indeed regulate cardiac muscle, then they do not appear to do so during development or young adulthood.

### *Gdf11^Gdf8aa^, Gdf8^Gdf11aa^*, and *Gdf8^Gdf11MD^* mice exhibit normal regeneration of damaged muscle after cryoinjury

It has been reported that muscle repair after toxin-induced injury to skeletal myofibers is significantly enhanced in *Mstn*-null mice (Wagner et al., 2005; McCroskery et al., 2005), suggesting that endogenous GDF8 suppresses satellite cell proliferation. Debate continues as to whether GDF8 acts directly on muscle satellite cells and whether such action may account, at least in part, for the muscle hyperplasia or hypertrophy observed in *Mstn*-null mice (Garikipati & Rodgers, 2012; George et al., 2013; Taylor et al., 2001; Thomas et al., 2000; Amthor et al., 2009; Lee et al., 2012; Walker et al., 2016). To test what impact enhancing GDF8 potency has on muscle regenerative capacity, we subjected *Gdf8^Gdf11aa^* and *Gdf8^Gdf11MD^* mutants to cryoinjury (**Fig. 6**). Tibialis anterior (TA) muscles of chimeric mice at 10-14 weeks of age were cryoinjured on day 0, harvested at 7 and 14 days post-injury, and analyzed via H&E staining (**Figs. 6, S6)**. Following cryoinjury, previously quiescent satellite cells in the basal lamina of myofibers are activated, giving rise to proliferating myoblasts (Dumont et al., 2015), which further differentiate and fuse together to form myotubes. Newly regenerated fibers can be distinguished by their central nuclei, with larger-sized fibers at early time points in the repair process corresponding to more advanced muscle fiber regeneration (Sinha et al., 2014; Cosgrove et al., 2014; Mauro, 1961). We similarly challenged *Gdf11^Gdf8aa^* mutants in muscle regeneration assays, as conflicting reports have been published regarding the impact of changing levels of GDF11 on muscle repair (Egerman et al., 2015; Sinha et al., 2014).

**Figure 6.**
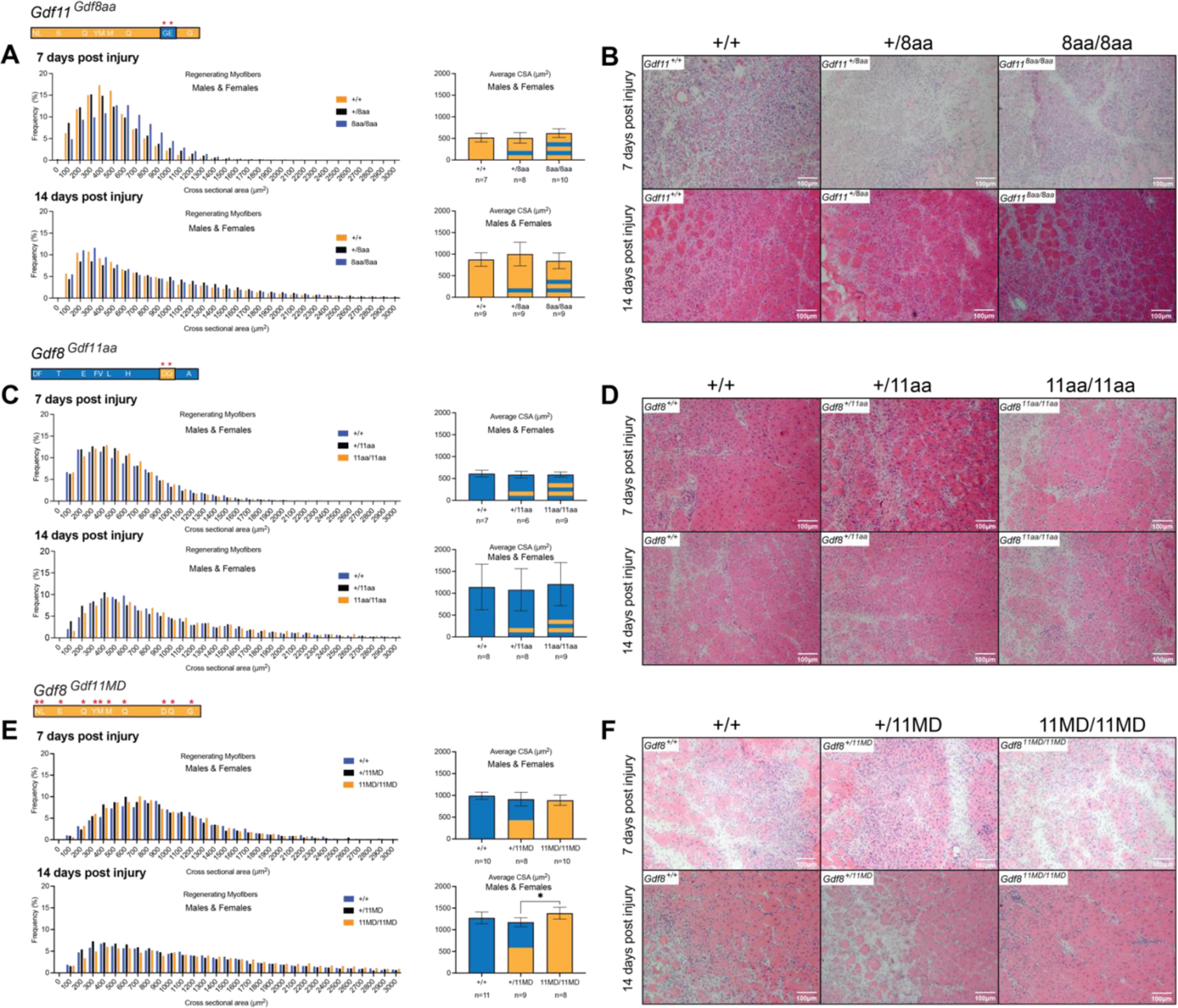
Regeneration in damaged muscle post cryoinjury in *Gdf11^Gdf8aa^, Gdf8^Gdf11aa^*, and *Gdf8^Gdf11MD^* mice. Tibialis anterior (TA) muscles were harvested from male and female mice at 7 and 14 days post-injury and analyzed by H&E staining **(B, D, F)**. The cross-sectional area (CSA) **(A, C, E)** of regenerating centrally nucleated fibers was measured using FIJI software (scale bar=100μm) and compared between genotypes across all three mouse lines at each timepoint. Overall, no significant differences were found in the size of regenerating myofibers in any genotype. However, at 14 days post-injury, bi-allelic *Gdf8^Gdf11MD/11MD^* mice exhibited modest increased cross-sectional area in comparison to *Gdf8^+/11MD^*, but not compared to *Gdf8^+/+^* mice. Statistical analysis was performed by one way ANOVA with Tukey’s correction for multiple comparisons. Also see Figure S6.

Analysis of regenerating myofibers in cryo-damaged muscles (7 or 14 days post-injury) showed no significant differences in average cross-sectional area (CSA) or fiber size distribution in either *Gdf8^Gdf11aa^* or *Gdf11^Gdf8aa^* mice (**Figs. 6A, B, C, D, S6)**. These data indicate that neither increasing the potency of GDF8 nor dampening that of GDF11 is sufficient to alter the time course or outcome of muscle fiber regeneration in young adult mice. Taken together with the lack of muscle or body weight change in *Gdf8^Gdf11aa^* or *Gdf11^Gdf8aa^* mice, it appears that postnatal skeletal muscle is more tolerant of changes in GDF11 signaling potency than embryonic bone, which exhibited skeletal transformations when GDF11 potency was reduced. In the *Gdf8^Gdf11MD^* mutants, regenerating fiber size in *Gdf8^+/11MD^* and *Gdf8^11MD/11MD^* mutants was indistinguishable from wild-type at both time points (**Fig. 6E, F)**. However, fiber size appeared slightly increased at 14 days post-injury in bi-allelic *Gdf8^11MD/11MD^* mutants compared to mono-allelic *Gdf8^+/11MD^* mutants (**Fig. 6E**). These data raise the possibility that the lack of mature GDF8, or the increased abundance of mature GDF11 protein, or the combination of these events might have a positive impact on the rate of muscle repair, consistent with prior reports of enhanced muscle regeneration following loss of GDF8 (Wagner et al., 2005; McCroskery et al., 2005) or supplementation of GDF11 *in vivo* (Sinha et al 2014).

Altogether, these data support the notion that the modest changes in ligand potency achieved through dual amino acid mutations confer a genetically-linked phenotype in early-stage skeletal development within only *Gdf11^Gdf8aa^* mutant mice. Additionally, the heightened ligand potency achieved through modifying the full mature domain of GDF8 in *Gdf8^Gdf11MD^* mutants confers a genetically-linked phenotype impacting skeletal muscle mass in adult mice. Incredibly, in bi-allelic *Gdf8^11MD/11MD^* mutants, circulating GDF11 concentration increased ∼50-fold more than normal, resulting in unexpectedly high levels of circulating GDF11 *in vivo*. Yet these chimeric mice were viable and survived normally into adulthood. The results indicate that gene regulation differences or differences in the prodomain regions may influence changes in ligand concentrations *in vivo*, and that ligand activity may not reflect alterations to ligand potency alone.

## Discussion

### Maintenance of GDF11 ligand potency and function is required for normal skeletal development

In this study, we investigated key phenotypes resulting from changes in the amino acid residues of GDF11 or GDF8, brought on by genetic modifications made to the *Gdf11* or *Gdf8* mature signaling domains *in vivo*. Results presented here indicate that changes in GDF11 and GDF8 potency elicit differential phenotypes (**Table 2**). Initial screening in early development revealed abnormal embryonic skeletal and vertebral transformations in the *Gdf11^8aa/8aa^* mutant mice, similar to those of mice with heterozygous (*Gdf11^+/-^*) deletion of *Gdf11* (McPherron et al., 1999). We followed these bi-allelic mutants of the *Gdf11^Gdf8aa^* line into adulthood to determine whether dampened GDF11 function in *Gdf11^Gdf8aa^* mutants results in additional phenotypic differences in organ growth and function postnatally. Assessments of overall body condition, skeletal and cardiac muscle mass, serum protein concentration, skeletal muscle repair, and baseline heart physiology and function were performed. Similar studies of the *Gdf8^Gdf11aa^* mutants were done in parallel to determine whether increased GDF8 potency in these mice resulted in differential phenotypes in adulthood. These efforts focused particularly on muscle size and function, given GDF8’s well-established negative regulation of muscle growth (Walker et al., 2016; McPherron et al., 1997; McPherron & Lee, 1997).

**Table 2.** Developmental patterns and phenotypes of *Gdf11^Gdf8aa^, Gdf8^Gdf11aa^*, and *Gdf8^Gdf11MD^* mice. Summary of phenotypic outcomes observed in *Gdf11^Gdf8aa^, Gdf8^Gdf11aa^*, and *Gdf8^Gdf11MD^* mice. Skeletal transformations were observed only in *Gdf11^Gdf8aa^* mice, resulting in the addition of one thoracic vertebra (T14 total) and vertebrosternal rib (T8 total) in bi-allelic *Gdf11^8aa/8aa^* mutants. In *Gdf8^Gdf11MD^* mice, bi-allelic *Gdf8^11MD/11MD^* mutants had increased levels of circulating GDF11 ∼50-fold higher than wild-type mice. *Gdf8^Gdf11MD^* mutants also presented with statistically significant decrease in skeletal muscle weights at 10-14 weeks of age. No mouse lines exhibited premature lethality or showed differences in overall heart physiology and function, kidney agenesis, liver weight, or muscle regeneration 7 and 14 days post-injury.

| Tissue / Phenotype | GDF11 | GDF8 |  |
| --- | --- | --- | --- |
| Predominant Expression Pattern | Developing limb buds; Primitive streak and tail bud | Development and adult skeletal muscle maintenance |  |
| **Chimeric Lines** | ***Gdf11<sup>Gdf8aa</sup>*** | ***Gdf8<sup>Gdf11aa</sup>*** | ***Gdf8<sup>Gdf11MD</sup>*** |
| Premature lethality | No | No | No |
| Bone | **Transformation of the axial skeleton (T14 thoracic vertebrae, T8 vertebrosteral ribs)** | No difference compared to wild-type | No difference compared to wild-type |
| Circulating ligand concentration | No difference compared to wild-type | No difference compared to wild-type | **~50-fold increase in GDF11 in bi-allelic mutants; GDF8 levels at or below level of detection** |
| Adult skeletal muscle | No difference compared to wild-type | No difference compared to wild-type | **Statistically significant decrease in mutant tibialis anterior, quadriceps, and triceps weights** |
| Heart | **Statistically significant increase in mutant male heart weights; not observed in mutant females** | No difference in function and physiology compared to wild-type | No difference in function and physiology compared to wild-type |
| Cardiac myocytes | No difference in cross-sectional area compared to wild-type | No difference in cross-sectional area compared to wild-type | No difference in cross-sectional area compared to wild-type |
| Kidney | Normal compared to wild-type; no observed renal agenesis | Normal compared to wild-type; no observed renal agenesis | Normal compared to wild-type; no observed renal agenesis |
| Liver | Observed slight increase in bi-allelic mutant liver weight, however not statistically significant | No difference compared to wild-type | **Statistically significant decrease in bi-allelic mutant female liver weight; not observed in mutant males** |
| Injured muscle regeneration | No difference compared to wild-type | No difference compared to wild-type | No difference compared to wild-type |

Our findings show that reducing the potency of mature GDF11 toward that of mature GDF8 is insufficient to sustain normal developmental function. Specifically, exchanging two amino acid residues from the *Gdf8* mature domain (G89 and E91) into the analogous location in the *Gdf11* mature domain (D89 and Q91) decreased the potency of mature GDF11 to that of GDF8 (Walker et al., 2017) and resulted in skeletal transformations detectable during early development in *Gdf11^Gdf8aa^* mice. However, exchanging the same amino acids from the *Gdf11* locus into the corresponding location in *Gdf8* did not produce similar physiological defects, and *Gdf8^Gdf11aa^* mice carrying one or both chimeric alleles were indistinguishable from wild-type mice in all of the assays we performed.

Because GDF8 and GDF11 both have been shown to play critical roles in skeletal and cardiac muscle development and function, and to regulate other organ systems (Walker et al., 2016; McPherron et al., 1997; Lee & Lee, 2013), further experiments were conducted to investigate the phenotypes of *Gdf11^Gdf8aa^* and *Gdf8^Gdf11aa^* mutant mice in early adulthood. Despite the axial skeletal defects found in the *Gdf11^Gdf8aa^* line, reduction in GDF11 potency in these mutants did not impact postnatal skeletal muscle growth or regenerative activity, nor did it alter baseline cardiac physiology or function into adulthood. We observed no differences in body weight or skeletal muscle weight (TA, quadriceps, and triceps muscle; normalized to body weight and tibia length) postnatally in male or female mutants at 10-14 weeks of age. Likewise, *Gdf8^Gdf11aa^* mice did not show significant differences in either sex at 10-14 weeks of age. For both chimeric lines, we saw no differences in heart or kidney weight in either the *Gdf11^Gdf8aa^* or *Gdf8^Gdf11aa^* mutants. The only statistically significant finding occurred in the heart weight of *Gdf11^8aa/8aa^* mutant males, compared to *Gdf11^+/+^* and *Gdf11^+/8aa^* mice; however, this result was not observed in mutant females of the same line. The differences in the heart weight normalized to body weight also proved insignificant in the *Gdf11^8aa/8aa^* mutants. Interestingly, *Gdf8^Gdf11aa^* mutants did not exhibit physiological changes to those found in *Gdf11^Gdf8aa^* mutants during embryonic development, nor did they produce measurable anatomic differences into adulthood.

### Replacement of GDF8 mature domain with GDF11 decreases skeletal muscle mass and produces a 50-fold increase of circulating GDF11 levels in young adult mutants

Similar to *Gdf8^Gdf11aa^* mutants, chimeric *Gdf8^Gdf11MD^* mice also did not exhibit detectable transformations during embryonic skeletal development, either in mono-allelic *Gdf8^+/11MD^* or bi-allelic *Gdf8^11MD/11MD^* mutants. Furthermore, we detected neither cranial bone malformation nor cleft palates in the mutants. Therefore, raising the activity of GDF8 to the full potency of GDF11 did not produce measurable changes in osteogenesis. This result may reflect the different tissue-specific expression patterns of the *Gdf11* and *Gdf8* loci. We performed additional experiments to determine whether increased GDF8 function in *Gdf8^Gdf11MD^* mutants might result in phenotypic differences in muscle growth and function postnatally and into adulthood. In *Gdf8^Gdf11MD^* mice, we confirmed that the increase in the GDF8 potency produced by replacing the mature domain of *Gdf8* with that of *Gdf11* decreased skeletal muscle size/mass in several limb muscles, consistent with results from GDF8 supplementation studies (Zimmers et al., 2002; Stolz et al., 2008) and with the recently reported *Mstn^Gdf11/Gdf11^* mice (Lee et al., 2022). Specifically, *Gdf8^Gdf11MD^* mutants exhibited decreased weight of the tibialis anterior (TA) and triceps muscles in early postnatal life in both male and female mice. A significant decrease in quadriceps muscle mass was also noted in mutant males, with bi-allelic mutants exhibiting the greatest change.

Our *Gdf8^Gdf11MD^* mutants also presented a striking outcome in terms of circulating levels of GDF11, which were increased to ∼50-fold more than normal, similar to the 30-40-fold increase in circulating GDF11 protein reported for *Mstn^Gdf11/Gdf11^* mice (Lee et al., 2022). While some prior studies suggested that elevation of GDF11— even at moderate levels—could result in detrimental consequences in mice, including severe cachexia and premature death (Harper et al., 2018; Egerman et al., 2015; Glass, 2016; Schafer et al., 2016), these results demonstrate that substantial elevation of GDF11 is well-tolerated, with only minor effects on skeletal muscle mass. *Gdf8^Gdf11MD^* chimeras were viable into adulthood and showed no signs of premature aging or other negative impacts on health or survival. The profound differences in circulating ligand levels in these gene-modified mice additionally suggest that gene regulatory differences encoded within the *Gdf8* and *Gdf11* genomic loci play an important role in determining systemic ligand abundance. Whether the substantial increase in GDF11 levels seen in *Gdf8^Gdf11MD^* mutants shifts the homeostasis of known antagonists or alters interactions with their respective N-terminal prodomains remains to be investigated. Finally, this study provides *in vivo* evidence supporting the molecular explanation for potency differences between GDF11 and GDF8 derived previously from structural and biochemical studies (Walker et al., 2017). In particular, while amino acid differences at residues 89 and 91 are responsible for much of the difference in potency observed between GDF11 and GDF8 (Walker et al., 2017), the other nine amino acids that distinguish the GDF11 and GDF8 mature domains clearly contribute as well.

We found no significant differences in heart weight or baseline cardiac physiology and function in *Gdf8^Gdf11MD^* mutants. However, whether similar results would be obtained in aged mutants or under conditions of transverse aortic constriction (TAC) remains a question for future studies. Although supra-physiologic elevation of GDF11 has been reported in several studies to impede recovery from muscle injury in young mice (Egerman et al., 2015; Hammers et al., 2017; Jones et al., 2018), and loss of GDF11 signaling in older animals has been suggested to underlie poorer regenerative outcomes with aging (Sinha et al., 2014), we also saw no difference in either the kinetics or ultimate outcome of muscle repair after injury in *Gdf8^Gdf11aa^* or *Gdf11^Gdf8aa^* mutants. With these data, we show that robust muscle repair activity is preserved in young mice both when GDF11 signaling is dampened and when GDF11 levels are raised, suggesting that muscle satellite cells and myofibers may be buffered to some extent against changes in GDF11 activity in young adulthood. The only other statistically significant finding occurred in the liver weight of *Gdf8^11MD/11MD^* mutant females, compared to *Gdf8^+/+^* mice and *Gdf8^+/11MD^* mutants, but not in *Gdf8^11MD/11MD^* mutant males. At this time, we cannot differentiate whether increased GDF8 potency directly affected liver hepatocytes, leading to the observed change in organ size, or whether indirect effects, potentially linked to changes in skeletal muscle or other tissues, underlie this result. Gene expression analysis in the liver to examine whether regulation of metabolic genes is altered will be useful to verify possible impact of the change in ligand potency.

Our chimeric mice, genetically modified to change unique amino acid residues between GDF11 and GDF8, demonstrate that sequence-determined structural differences in these ligands are critically important, and not simply accounted for by gene regulation differences alone. We have discovered that two specific amino acids in the fingertip region of the GDF11 mature domain are required for proper axial skeletal patterning during early-stage development, but do not appear to be crucial for regulating heart or skeletal muscle. Additionally, substituting the full mature domain sequence of GDF11 into the *Gdf8* locus, in place of mature GDF8, created a GDF8-null mouse with decreased skeletal muscle mass. Circulating GDF11 concentrations in this mutant were significantly higher than wild-type, showing that very high GDF11 blood levels can be tolerated and can overcome phenotypes typically associated with loss of GDF8 function. Our findings show that changing the ligand potency of GDF11 and GDF8 or altering their bioavailability within the mammalian system causes distinct measurable physiological effects, and that maintenance of functional GDF11 and GDF8 are necessary for proper development and adult tissue maintenance.

## Materials & Methods

### Lead Contact

For original materials and data, please contact Richard T. Lee at. No new or unpublished custom code, software, or algorithm was generated for this work.

### Mice caretaking

Mice handling and experimentation followed guidelines set forth by the Institutional Animal Care and Use Committee (IACUC) at Harvard University/Faculty of Arts and Sciences. Regular animal housing and care were carried out by the Biological Laboratories staff in accordance with relevant IACUC regulations and guidelines. Housing included density of 2-4 mice per cage, along with Enviro-Dri bedding, cotton nestlet, one red hut, automatic waterspout dispensing reverse osmosis deionized water, and regular chow diet (Prolab IsoPro RMG3000 5P75/76).

### Generation of mutant mouse lines

The CRISPR/Cas9 system was used to generate chimeric mice carrying amino acid mutations from the GDF8 and GDF11 interchanged in the mature domain. Two rounds of microinjections into C57BL/6J zygotes were performed for each line, delivering (1.) purified *Streptococcus pyogenes* Cas9 (spCas9) mRNA, (2.) *in vitro*-transcribed synthetic guide RNAs (sgRNA), which targeted the native loci near the desired integration sites, and (3.) the generated *Gdf11* and *Gdf8* ssDNA or *Gdf11* dsDNA donor template. Zygote injections were performed by the Genome Modification Facility at Harvard University. Super-ovulated C57BL/6J female mice were mated to C57BL/6J males, and the fertilized zygotes were subsequently harvested from the oviducts. Zygotic pronuclei were microinjected with (1) purified *spCas9* mRNA (System Bioscience, 100ng/μl), (2) *in vitro*-transcribed synthetic guide RNAs (sgRNA) which targeted near the integration sites of the native loci, and (3) the generated *Gdf11* and *Gdf8* ssDNA donor construct. Microinjected zygotes were implanted into the oviducts of C57BL/6J surrogate females at 12 hours post coitum. To identify mosaic offspring that contained the desired chimeric *Gdf11* and *Gdf8* double amino acid substituted sequences, progeny from the microinjected surrogate females were genotyped by Sanger sequencing and by PCR validation and subcloning. In the *Gdf11* amino acid chimera, we produced 47 total pups, with 14 live pups and 33 dead pups. Of the 14 live pups, 4 exhibited a mosaic genotype, and 3 were ultimately selected as germline founders. In the *Gdf8* amino acid chimera, we obtained 92 viable pups, with 0 dead pups. Of the 92 live pups, 49 showed a mosaic genotype upon screening for the targeted allele. Ultimately, 4 confirmed positive founders were used as breeders to establish the colony. In the *Gdf8* full mature domain chimera, we obtained 80 viable pups, with 2 dead pups. Of the 80 live pups, 35 showed a mosaic genotype upon screening for the targeted allele. Ultimately, 5 confirmed positive founders were used as breeders to establish the colony. We bred these chosen mice with wild-type C57BL/6J mice and genotyped the resultant pups to confirm that our desired mutation was incorporated in the germline. We backcrossed the knock-in alleles 5 generations in a C57BL/6J genetic background prior to characterization experiments. Male and female mice were selected based on gender and randomized prior to treatment for all proposed animal studies.

### Genotyping

Initial mouse colony breeder genotypes were verified by sequencing. Subsequent progeny tissues were collected from the tail or the ear and mixed in 1.5ml Eppendorf tubes containing 300μl of lysis solution (50mM KCl, 10mM Tris–HCl pH8.3, 2.5mM MgCl_2_, 0.1mg/ml gelatin, 0.45%NP40, and 0.45% Tween-20 in ddH_2_O) with 10μg/ml Proteinase K and incubated at 50°C for 10-12 hours. Next, samples were placed in heating blocks at 98°C for 5 minutes to inactivate the Proteinase K. DNA was extracted using a standard phenol-chloroform/ethanol precipitation protocol, and genotyping was performed by PCR using Phusion High-Fidelity PCR Master Mix with HF Buffer. Primers used are as follows: For *Gdf11^Gdf8aa^*, FW: CCTGACCCTCAGCATCCTTTCA, RV: GGTCCTTACTTTGCCCCATCCT; for *Gdf8^Gdf11aa^* and *Gdf8^Gdf11MD^*, FW: TGTGGTTGGTTTGTTTGTTTGT, RV: GCCTGTGGTGCTTGAATTCA. PCR products were digested with restriction enzyme *AseI* (NEB R0526), with NEBuffer 3.1, at 37°C for 20-30 minutes and analyzed on 1% agarose gel. Further genotyping was performed via Sanger sequencing to verify the nucleotide changes and presence of the *AseI* site.

### Targeted locus amplification (TLA) sequencing

Bone marrow from *Gdf11^Gdf8aa^, Gdf8^Gdf11aa^*, and *Gdf8^Gdf11MD^* F5 generation mutant mice were harvested, homogenized, and subjected to ACK lysis. Harvested bone marrow (5 vials, each containing 1 x 10^7^ cells) was frozen and delivered to Cergentis B.V. (Utrecht, the Netherlands) for TLA sequencing analysis (de Vree et al., 2014). TLA sequencing used a locus-specific sequence for the targeted amplification and complete sequencing of *Gdf11* and *Gdf8* loci. The genomic DNA was crosslinked, digested and re-ligated, before it was purified and circular TLA fragments were then amplified with two independent sets of inverse primers, corresponding to the *Gdf11* or *Gdf8* locus specific transgene, to identify the location of each targeting event across the whole genome. Primer sets used are as follows.

*Gdf11^Gdf8aa^*: Upstream, Fw:ACATTTGCTCCCATTACTGT, Rv:AGCAATAAGAACAAGGGAGC, Downstream, Fw:CAAGAGTCTTAAGAGGATGGG, Rv:GGGTAGTTTAGTAGCTCTCATAG.

*Gdf8^Gdf11aa^*: Upstream, Fw:GAATAGATGCAATGGTTGGC, Rv:AGAGTGTAGTGTTTAAGTAGCA, Downstream, Fw:CACAATTTGTTTATGCGGTTT, Rv:TCTCACTTCCTTGCCTAGAT.

*Gdf8^Gdf11MD^*: chr10 detection, Fw:CAAGTGGGTGTGTGGATAC, Rv:CTACCAAGATGTCCCCAATC, chr1 detection, Fw: GTAACTGCTCAGATTCCCAA, Rv: AGCTATTCCAAGGAACAACA. 5’ integration site: chr1:53,066,297 (tail) fused to Insert: 1, head (the same as chr10:128,885,435, tail) with 4 inserted bases ATCCCTTTTTAGAAGTCAAGGTGACAGACACACCCAAGAGGTCCCGAAGAAACCTAGGCCTGGACTGGATGAACACTCGAGTGAGTCCCGCTGCTGCCGATATCCTCTCACAGTGGACTTTGAGGCTTTTGGCTGGACTGGATCATC; 3’ integration site: Insert: 327, tail (the same as chr10:128,885,108, head) fused to chr1:53,066,629 (head) AACATGCTCTACTTCAATGACAAGCAGCAGATTATCTACGGCAAGATCCCTGGCATGGTGGTGGATCGATGTGGCTGCTCCTGAGCTTTGCATTAGGTTAGAAATTTTCCAAGTCATGGAAGGT CTTC. Following amplification of the targeted locus, PCR amplicons of the complete *Gdf11* and *Gdf8* region of interest were purified and prepped for Illumina sequencing. Analysis was performed by aligning mutant sequences to the mouse mm10 reference genome sequence. In *Gdf11^Gdf8aa^* mice, two mutated nucleotides (G→C and T→C) were confirmed on chr10 in the mature domain of *Gdf11*, resulting in alteration of only the two targeted amino acids: *Gdf11* D89 to G89 (Asp→Gly) and Q91 to E91 (Gln→Glu). In *Gdf8^Gdf11aa^* mice, two mutated nucleotides (G→A and G→C) were confirmed on chr1 in the mature domain of *Gdf8*, resulting in amino acid changes in *Gdf8* G89 to D89 (Gly→Asp) and E91 to Q91 (Glu→Gln). In bi-allelic *Gdf11^8aa/8aa^* and *Gdf8^11aa/11aa^* samples, the mutated nucleotides were confirmed at 100% frequency, indicating that the mutations occurred on both alleles, while in mono-allelic *Gdf11^+/8aa^* and *Gdf8^+/11aa^* samples, the mutations were confirmed at ∼50% frequency, indicating occurrence of the desired mutations on only one allele. In the *Gdf8^Gdf11MD^* mice, correct integration of exon 3 of *Gdf11* was confirmed on chr1 in place of the native exon 3 of *Gdf8*, indicating successful replacement of the GDF8 mature domain with that of GDF11. In bi-allelic *Gdf8^11MD/11MD^* samples, no wild-type reads were present at the integration site and in the deleted region, confirming bi-allelic replacement of native GDF8, while in mono-allelic *Gdf8^+/11MD^* samples, wild-type reads were detected at the integration site and in the deleted region on one allele, confirming mono-allelic replacement of native GDF8.

### Serum mass spectrometry of GDF11 and GDF8

For serum collection, after euthanasia, blood was collected via orbital bleeding into Microtainer tubes with a serum separator (BD) and incubated for 30 minutes at room temperature before centrifugation at 2000xG for 10 minutes at room temperature. The upper layer of serum was then transferred to a new tube and stored at −80°C prior to mass spectrometry analysis. Serum from chimeric mice (minimum 100μl) was submitted to the Brigham and Women’s Hospital Brigham Research Assay Core (BRAC) for quantitative liquid chromatography tandem mass spectrometry detection of GDF11 and GDF8 protein concentrations. The mouse serum was denatured and alkylated, followed by pH-based fractionation, using cation ion exchange SPE. After desalting and concentrating, the peptide mix was separated via liquid chromatography, followed by mass spectrometry analysis in a positive electrospray ionization mode. GDF11 and GDF8 concentrations were determined using unique proteotypic peptides from GDF11 and GDF8 as surrogate peptides, coupled with heavy-labeled unique peptides as internal standards. Included in the analysis were GDF8 concentrations and the mean GDF11 concentrations.

### Skeletal preparation

Embryonic day 18.5 (E18.5) harvests (Lewandowski et al., 2019) were performed for skeletal and vertebral analyses of *Gdf11^Gdf8aa^* and *Gdf8^Gdf11aa^* mutants and compared with wild-type mice. For timed breeding, a vaginal plug observed in the female indicated embryonic day 0.5 (E0.5). Mouse embryos were harvested at E18.5, skinned, eviscerated, and underwent washes of 100% ethanol and 100% acetone for 24 hours at room temperature. The skeletons were then stained using a 0.3% alcian blue (dyes bone) and 0.1% Alizarin red (dyes cartilage) solution at 37°C while oscillating for 72 hours. Next, they were transferred to 1% KOH for 24 hours on a rocker at room temperature, followed by a series of glycerol/KOH washes at (1) 20% glycerol/1% KOH, (2) 50% glycerol/1% KOH, and (3) 80% glycerol/1% KOH for 24 hours at room temperature. The stained preparations were placed in 80% glycerol/1XPBS and imaged using a Nikon D750 camera attached to a Nikon SMZ1500 stereo microscope with an HR Plan APO 1x objective lens. Images were captured in NEF (Nikon Electronic Format) and processed and adjusted in the Adobe Camera Raw platform.

### Tissue collection and analysis

Mice were ear-tagged and their genotype blinded at 10-14 weeks of age. Adult mice were euthanized with CO_2_, and overall body weights were taken postmortem. For tissue collection, the heart was dissected, washed in PBS, and dried on paper towels prior to weighing. Th e tibialis anterior muscle, quadriceps muscle, and triceps muscle were harvested and weighed. The tibia bone was excised, cleaned, and measured using electronic calipers. Statistical analysis was performed using Prism 8.4.2 for macOS. The muscle weights were divided by sex and genotype within each mouse line, and the results were represented as the mean within each group by genotype and sex. The combined muscle weights were normalized to body weight and tibia length. Groups more than two were compared by one way ANOVA with Tukey’s correction for multiple comparisons.

### Echocardiographic studies

*Gdf11^Gdf8aa^* mice (*Gdf11^+/+^*, n=12; *Gdf11^+/8aa^*, n=16; and *Gdf11^8aa/8aa^*, n=15) and *Gdf8^Gdf11aa^* mice (*Gdf8^+/+^*, n=9; *Gdf8^+/11aa^*, n=10; and *Gdf8^11aa/11aa^*, n=10), age 10-14 weeks, were sedated with 0.1–0.5% inhaled isoflurane for echocardiography (Loffredo et al., 2013). Mice were placed on a heating pad, and echocardiograms were obtained at mid-papillary level with the Vevo3100 (Visualsonics, Canada). The heart rate of every mouse studied was monitored and maintained at >400bpm during echocardiographic procedure, so as to (1) avoid artificial myocardial depression brought on by exposure to isoflurane during imaging and (2) ensure consistent measurements across all study groups. Parasternal long-axis views, short-axis views, and two-dimensional M-mode was used to measure left ventricular interventricular septal wall thickness (IVS/LVAW), left ventricular posterior wall thickness (LVPW), and left ventricular internal diameter (LVID) during both systole and diastole in both mouse lines. Fractional shortening (FS,%) was calculated with the Visualsonics software package. Statistical analysis was performed using Prism 8.4.2 for macOS. For echocardiographic and morphometric analyses, cross sectional area (CSA), capillary density and volume results were presented as the mean for each group by genotype and sex. At least three measurements were averaged and used for every data point from each mouse. Groups more than two were compared by one way ANOVA with Tukey’s correction for multiple comparisons. All analyses were performed under blinded conditions.

### Muscle cryoinjury

The muscle cryoinjury procedure (Oh et al., 2016) was chosen due to its ability to generate a reproducible injury area with a discrete border between injured and uninjured muscle. This border remains distinct during regeneration of injured muscle. Mice were anesthetized using isoflurane and the skin over the left tibialis anterior (TA) muscle was shaved and disinfected using Betadine, followed by wiping with 70% ethanol. The TA was then exposed by a small 3mm incision. A metal probe with flat round bottom cooled down in dry ice was applied directly to exposed TA muscle for 5 seconds. The skin incision was then closed with synthetic absorbable suture (5-0 coated vicryl) immediately following the injury. Buprenorphine (0.05-0.1mg/kg, s.c.) was administered immediately after recovery from surgery, and subsequently every 8-12 hours, for at least 48 hours after surgery. Injured muscles were recovered for 7- or 14-days post injury. The cryoinjury model employed has been widely utilized to assess muscle repair after damage and offers a number of advantages. In particular, cryoinjury can be performed such that the size of the lesion and severity of damage is highly similar across experimental animals, with preservation of regenerative muscle satellite cells that nucleate repair in the surrounding uninjured area (Dumont et al., 2015; Gayraud-Morel, 2009; Hardy et al., 2016).

### Histology and cross-sectional area quantification

For cryosections, harvested TA muscles were flash frozen in 2-methylbutane for 30 seconds followed by liquid nitrogen for 30 seconds. Samples were stored at −80°C prior to sectioning. Mouse hearts were rinsed with 1x PBS, embedded in compound at optimal cutting temperature and frozen. TA samples and cardiomyocyte samples were sectioned at 10μm. Hematoxylin and eosin (H&E) staining was used to visualize cardiomyocyte cross-sections and regenerating myofibers in injured muscles. Sections were stained with Hematoxylin for 3 minutes, washed with tap water for 2 minutes, followed by back-to-back washes in Acid Alcohol. Scott’s Bluing Reagent was used for 3 minutes for nuclear staining, followed by another tap water wash, and finally, 2 minutes of eosin staining. Afterward, sections were dehydrated in ethanol and xylene, and subsequently mounted in Permount Mounting Medium (catalog 17986-01, Electron Microscopy Sciences) and cured for 24 hours before imaging. In regenerating myofibers after cryoinjury, a cross section representing the mid-belly of the TA where there was a clear representation of the injury was chosen. To quantify muscle fiber size, a series of images were taken spanning the entire regenerating area in the cross sections using a dual head Olympus BX4 microscope with cellSens Standard software. Centrally nucleated myofibers were measured in each image using FIJI software (scale bar=100μm), resulting in ∼300-1500 fibers collectively for each animal. Comparative analyses of more than two groups were performed by one way ANOVA with Tukey’s correction for multiple comparisons. All analyses were performed under conditions in which the analyst was blinded to sample identity.

### Statistical Analyses

All data are presented as mean + standard error of mean (s. e. m.). For all data, *n* equals the number of biological replicates of animals used per experiment. The number of animals used for each group was determined based on total empirical data and anticipated completeness of datasets and was sufficient to detect differences in experimental outcomes, if present. Statistical analysis was performed using Prism 8.4.2 for macOS. Comparisons between two different experimental groups were assessed for statistical significance using Student’s *t* test, and comparisons between more than two groups were compared by one way ANOVA with Tukey’s correction for multiple comparisons. Statistical significance was accepted at p<0.05. All experiments and analyses were performed under blinded conditions.

## Acknowledgements

We thank Lin Wu and Laurie Chen from Harvard’s Genome Modification Facility for assistance in generating the *Gdf11* and *Gdf8* chimeric mice, Liming Peng and Shalender Bhasin for assistance with tandem liquid-chromatography/mass spectrometry, Cergentis B.V. for TLA sequencing analysis, Thompson and Wagers lab members for helpful discussions.

## Author Contributions

J.L. and R.G.W. conceived and designed the study. J.L., R.G.W., A.D’A., A.V., K.A.M., M.J.M., K.R.M., and J.M. G. performed experiments and analyzed data. K.A.L. performed statistical analyses. S.E. and E.V.S. provided technical assistance. J.L., R.G.W., A.D’A., A.V., K.A.M., T.B.T., A.J.W., and R.T.L. performed formal data analysis and interpretation. J.L. and R.G.W. wrote the manuscript. All authors reviewed the manuscript.

