## Supplemental Figures and Tables for "Functional substitutions of amino acids that differ between GDF11 and GDF8 impact skeletal development and skeletal muscle"

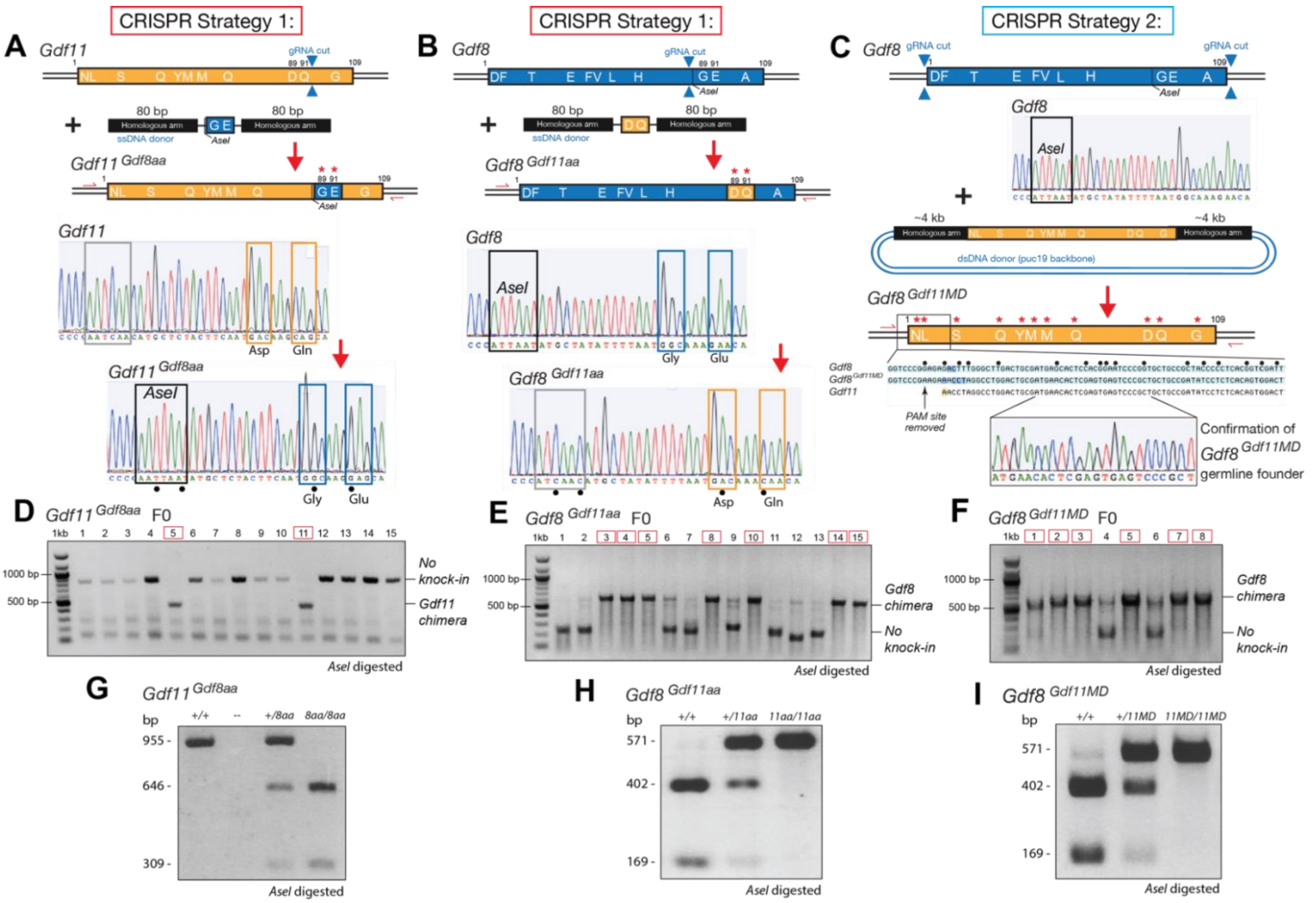


**Supplemental Figure 1. CRISPR/Cas9 approach and characterization for generating GDF11 and GDF8 chimeric mice.** To generate *Gdf11*$^{Gdf8aa}$, *Gdf8*$^{Gdf11aa}$, and *Gdf8*$^{Gdf11MD}$ mutant lines, DNA donor plasmids were designed to contain the desired amino acid or full mature domain mutations. For the amino acid mutant lines *Gdf11*$^{Gdf8aa}$ and *Gdf8*$^{Gdf11aa}$, single-stranded donor templates were flanked by 80 base homologous arms, while a double-stranded donor template was flanked by 4kb homologous arms was constructed for the full mature domain mutant *Gdf8*$^{Gdf11MD}$. Schematics of the CRISPR strategy depict the native *Gdf11* **(A)** and *Gdf8* **(B, C)** loci, with locations of the gRNA cut site (top), the donor templates containing the codons to incorporate the *Gdf8*-like and *Gdf11*-like amino acid mutations (middle), and the resulting chimeric mutants (bottom). Successful transgene modification was confirmed via Sanger sequencing **(A, B, C)** and genotyping by PCR validation and subcloning **(D, E, F)** were performed to identify F0 founders. **(G)** After genotyping, digested bands from incubation of DNA with *AseI* is shown for *Gdf11*$^{+/8aa}$ and *Gdf11*$^{8aa/8aa}$ samples, which acquired the *AseI* site from the donor template. **(H)** *Gdf8*$^{+/11aa}$ and *Gdf8*$^{11aa/11aa}$ DNA can no longer be digested fully after incubation with *AseI*, indicating absence of the excised native *AseI* site. **(I)** *Gdf8*$^{+/11MD}$ and *Gdf8*$^{11MD/11MD}$ DNA also cannot be digested fully after incubation with *AseI*. Amino acid residues of *Gdf11* represented in orange and *Gdf8* in blue.

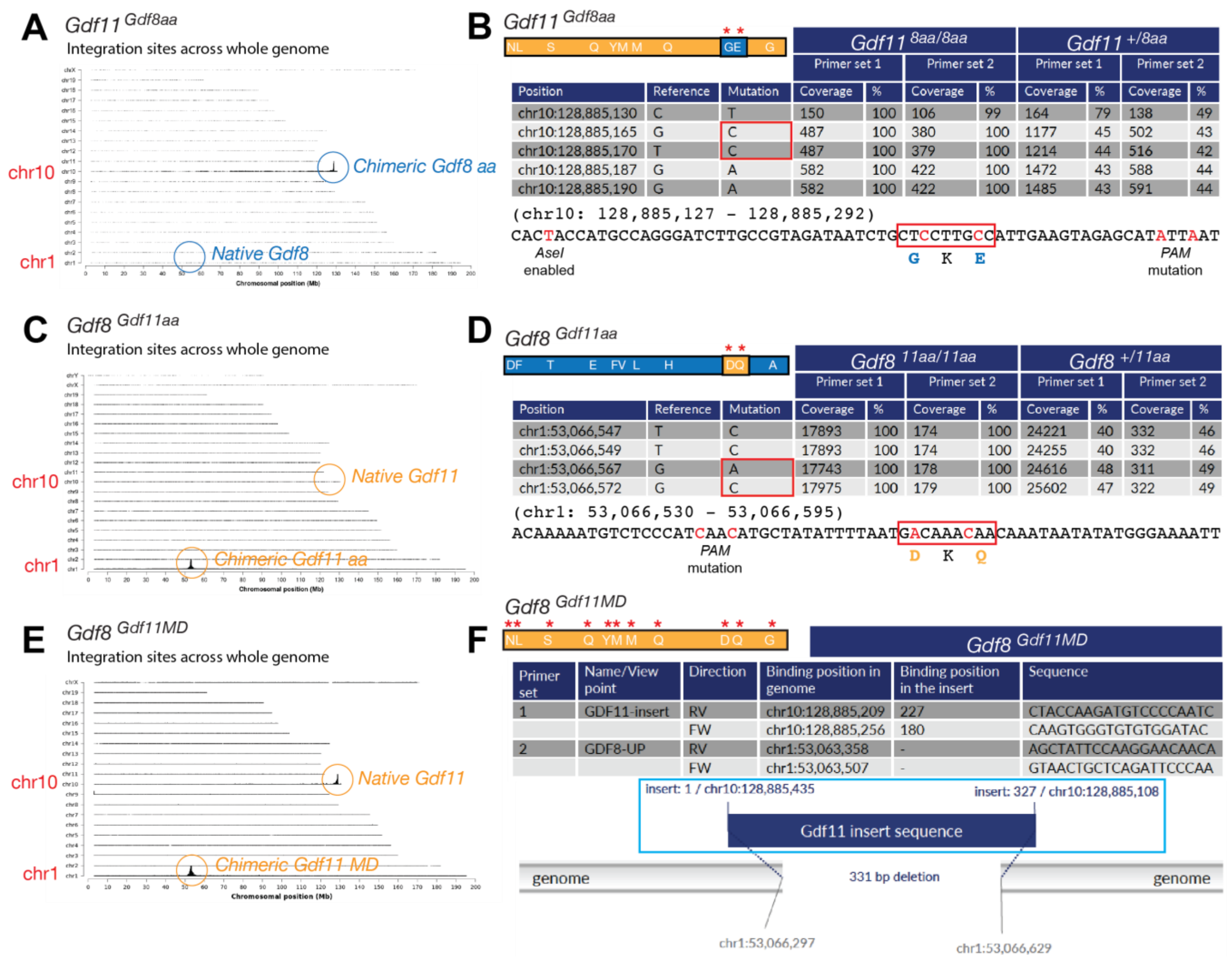


**Supplemental Figure 2. Genotyping and Targeted Locus Amplification (TLA) sequencing of whole genome integration events in *Gdf11*$^{\mathit{Gdf8aa}}$, *Gdf8*$^{\mathit{Gdf11aa}}$, and *Gdf8*$^{\mathit{Gdf11MD}}$ mutants.** In *Gdf11*$^{\mathit{Gdf8aa}}$ mice, detected mutations were in exon 3 of *Gdf11* on chr10 (RefSeq) **(A)**. In *Gdf11*$^{\mathit{8aa/8aa}}$ samples **(B)**, 2 mutated bases (G→C = D89G and T→C = Q91E) were found at 100% frequency, indicating bi-allelic mutation. In *Gdf11*$^{\mathit{+/8aa}}$ samples, the mutations occurred at ~50% frequency, indicating mono-allelic mutation. Three additional silent mutations **(B)**—one for the insertion of the *AseI* site unique to *Gdf8* and two for the mutations—were also confirmed. In *Gdf8*$^{\mathit{Gdf11aa}}$ mice, mutations were in exon 3 of *Gdf8* on chr1 (RefSeq) **(C)**. In *Gdf8*$^{\mathit{11aa/11aa}}$ samples **(D)**, 2 mutated bases (G→A = G89D and G→C = E91Q) were found at 100% frequency, indicating bi-allelic mutation. In *Gdf8*$^{\mathit{+/11aa}}$ samples, the mutations occurred at ~50% frequency, indicating mono-allelic mutation. Two additional base changes **(D)**, accounting for the PAM site silent mutation, and the absence of the *AseI* site were also confirmed. In *Gdf8*$^{\mathit{Gdf11MD}}$ mutants, the full GDF11 mature domain sequence was found on chr1 **(E)**, in addition to the native *Gdf8* locus on chr10. No structural variants, off-target integration, or locus duplication were detected. *Gdf8*$^{\mathit{11MD/11MD}}$ samples lacked wild-type *Gdf8* mature domain sequences at the integration site and in the deleted region, confirming bi-allelic replacement of native GDF8, while *Gdf8*$^{\mathit{+/11MD}}$ samples had wild-type reads at the integration site and in the deleted region on one allele, confirming mono-allelic replacement of native GDF8. The mouse mm10 reference sequence was used for comparisons. Schematic of the amino acid residues in *Gdf11* (orange) and *Gdf8* (blue) locus.

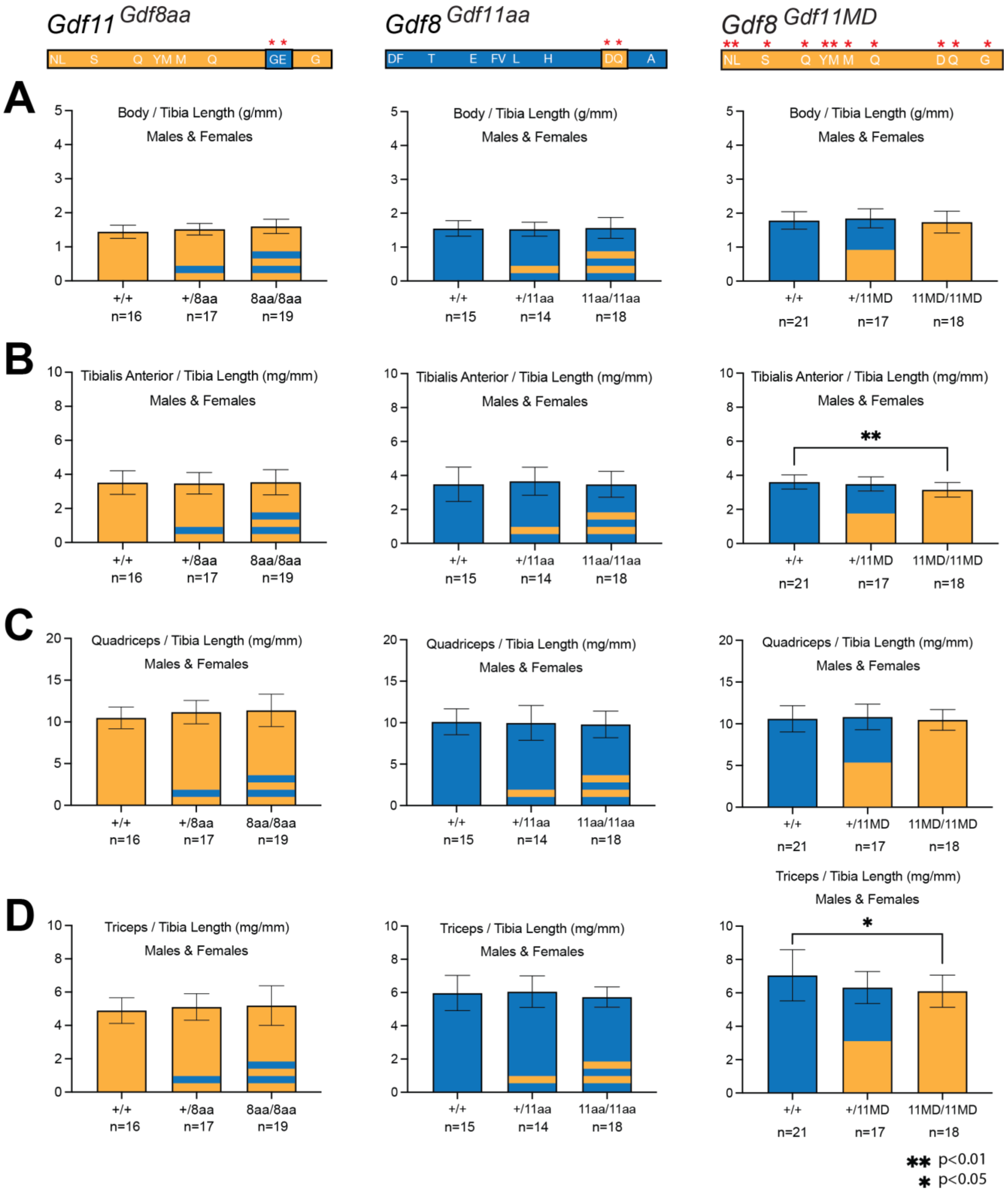


**Supplemental Figure 3. Skeletal muscle measurements normalized to tibia length for *Gdf11$^{Gdf8aa}$*, *Gdf8$^{Gdf11aa}$*, and *Gdf8$^{Gdf11MD}$* mice.** The overall body weight **(A)**, tibialis anterior TA muscle **(B)**, quadriceps muscle **(C)**, and triceps muscle **(D)** weights of *Gdf11$^{Gdf8aa}$*, *Gdf8$^{Gdf11aa}$*, and *Gdf8$^{Gdf11MD}$* mice at 10-14 weeks of age were normalized to tibia length. In *Gdf8$^{Gdf11MD}$* mice, there was a statistically significant decrease in the TA muscle **(B)** normalized to tibia length between *Gdf8$^{11MD/11MD}$* (n=18) vs. *Gdf8$^{+/+}$* (n=21, p<0.01) mice, and in the triceps muscle **(D)** normalized to tibia length between *Gdf8$^{11MD/11MD}$* (n=18) vs. *Gdf8$^{+/11MD}$* (n=21, p<0.05) mice. Statistical analysis was performed by one way ANOVA with Tukey's correction for multiple comparisons.

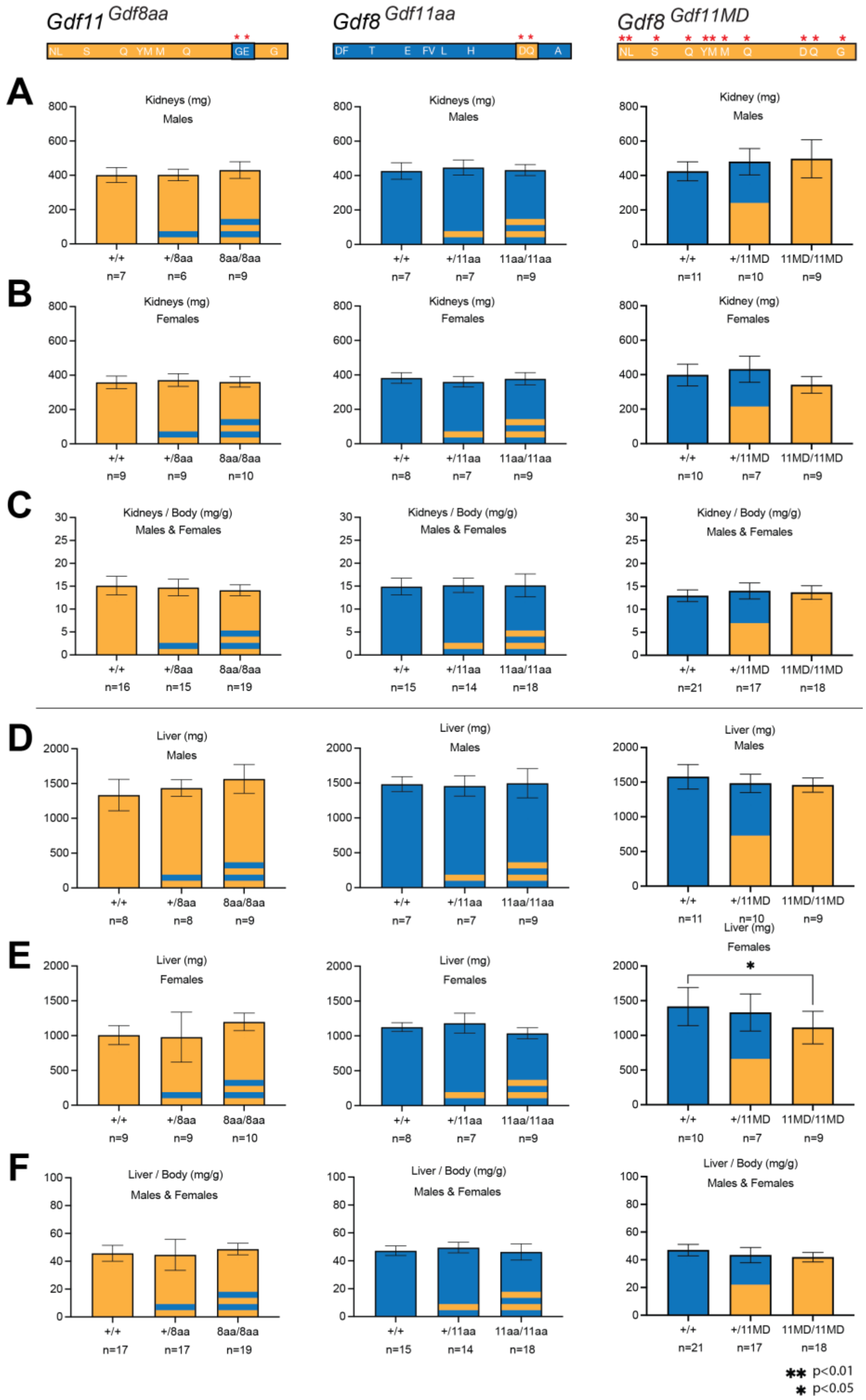


**Supplemental Figure 4. Kidney and liver weights for *Gdf11*$^{Gdf8aa}$, *Gdf8*$^{Gdf11aa}$ mice, and *Gdf8*$^{Gdf11MD}$ mice.** Kidney weight (of 2 kidneys) **(A, B)** and liver weight **(D, E)** were taken from *Gdf11*$^{Gdf8aa}$, *Gdf8*$^{Gdf11aa}$ mice, and *Gdf8*$^{Gdf11MD}$ mice at 10-14 weeks of age and normalized to overall body weight (C, F). No mutant mouse of any chimeric line exhibited fewer than two kidneys. In *Gdf11*$^{Gdf8aa}$, *Gdf8*$^{Gdf11aa}$ mice, no significant differences were found in mutant mice, compared to wild-type. A significant decrease was found in the liver weight of female *Gdf8*$^{11MD/11MD}$ mutants (n=9) **(E)**, compared to *Gdf8*$^{+/+}$ mice (n=10, $p<0.05$). However, the corresponding difference was not found in the liver weight of males **(D)** or the normalized weight of male and female *Gdf8*$^{Gdf11MD}$ mice **(F)**. Statistical analysis was performed by one way ANOVA with Tukey's correction for multiple comparisons.

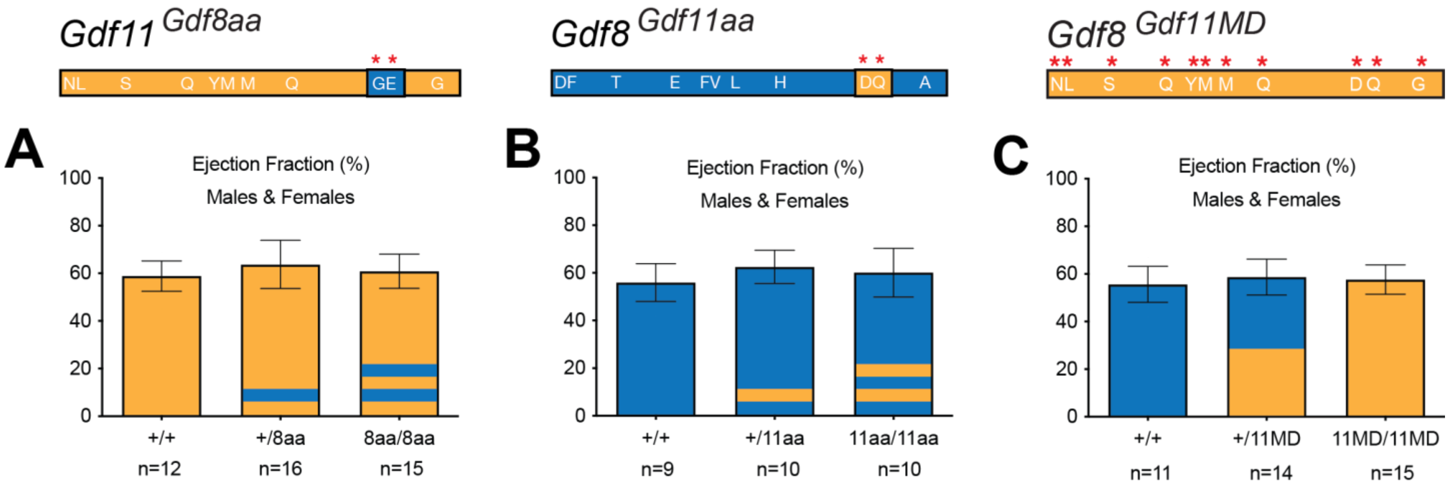


**Supplemental Figure 5. Cardiac ejection fraction readings of *Gdf11*$^{\textit{Gdf8aa}}$, *Gdf8*$^{\textit{Gdf11aa}}$, and *Gdf8*$^{\textit{Gdf11MD}}$ mice.** Ejection fraction (ES) measurements were taken and compared between genotypes of *Gdf11*$^{\textit{Gdf8aa}}$ mice **(A)**, *Gdf8*$^{\textit{Gdf11aa}}$ **(B)**, and *Gdf8*$^{\textit{Gdf11MD}}$ mice **(C)**, with no significant differences found. Statistical analysis was performed by one way ANOVA with Tukey's correction for multiple comparisons.

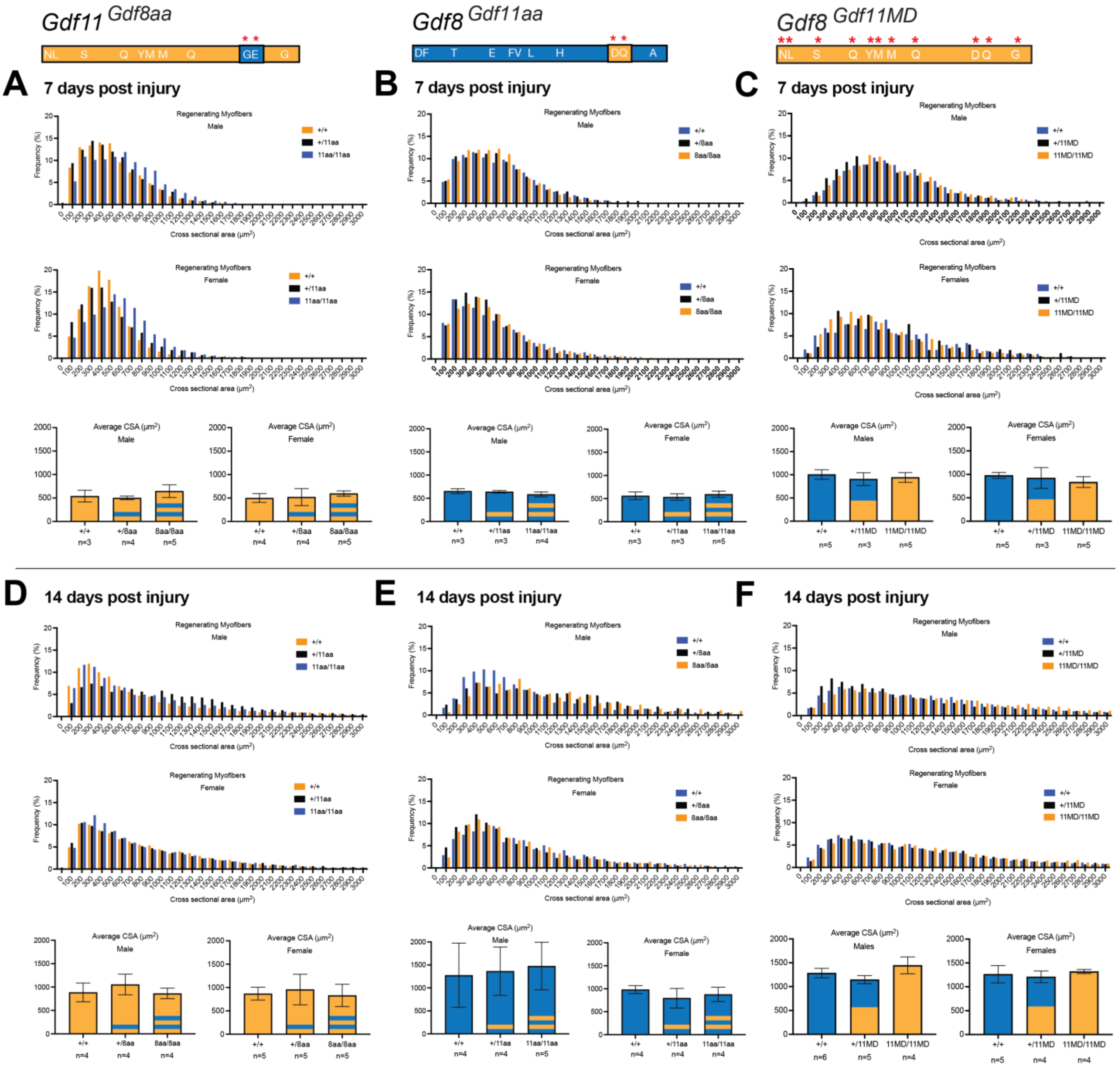


**Supplemental Figure 6. Normal regeneration of damaged muscle in male and female *Gdf11*$^{Gdf8aa}$, *Gdf8*$^{Gdf11aa}$, and *Gdf8*$^{Gdf11MD}$ mice after cryoinjury.** Tibialis anterior (TA) muscles of male and female mice were harvested at 7 days **(A, B, C)** and 14 days **(D, E, F)** post-injury and analyzed by H&E staining. The cross-sectional area (CSA) of regenerating centrally nucleated fibers was measured using FIJI software (scale bar=100μm). Statistical analysis was performed by one way ANOVA with Tukey's correction for multiple comparisons.

| Mouse Line | *$Gdf11^{Gdf8aa}$* | | | *$Gdf8^{Gdf11aa}$* | | | *$Gdf8^{Gdf11MD}$* | | |
|---|---|---|---|---|---|---|---|---|---|
| **Genotype** | **+/+** | **+/8aa** | **8aa/8aa** | **+/+** | **+/11aa** | **11aa/11aa** | **+/+** | **+/11MD** | **11MD/11MD** |
| **F4 Breeding Pairs** | **F5 Progeny** | | | **F5 Progeny** | | | **F5 Progeny** | | |
| **(Round 1)** | **(Round 1)** | | | **(Round 1)** | | | **(Round 1)** | | |
| +/X x +/X | 2 (20.0) | 5 (50.0) | 3 (30.0) | 3 (30.0) | 4 (40.0) | 3 (30.0) | 4 (25.0) | 8 (50.0) | 4 (25.0) |
| +/X x X/X | -- | 5 (55.6) | 4 (44.4) | -- | 4 (44.4) | 5 (55.6) | -- | 5 (55.6) | 4 (44.4) |
| X/X x X/X | -- | -- | 8 (100) | -- | -- | 10 (100) | -- | -- | 7 (100) |
| **(Round 2)** | **(Round 2)** | | | **(Round 2)** | | | **(Round 2)** | | |
| +/X x +/X | 3 (23.1) | 6 (46.2) | 4 (30.7) | 2 (18.2) | 5 (45.5) | 4 (36.3) | 3 (25.0) | 5 (41.7) | 4 (33.3) |
| +/X x X/X | -- | 5 (62.5) | 3 (37.5) | -- | 4 (50.0) | 4 (50.0) | -- | 4 (50.0) | 4 (50.0) |
| X/X x X/X | -- | -- | 9 (100) | -- | -- | 11 (100) | -- | -- | 11 (100) |
| **(Round 3)** | **(Round 3)** | | | **(Round 3)** | | | **(Round 3)** | | |
| +/X x +/X | 3 (30.0) | 5 (50.0) | 2 (20.0) | 3 (30.0) | 5 (50.0) | 2 (20.0) | 4 (21.1) | 10 (52.6) | 5 (26.3) |
| +/X x X/X | -- | 4 (40.0) | 6 (60.0) | -- | 6 (54.5) | 5 (45.5) | -- | 7 (58.3.0) | 5 (41.7) |
| X/X x X/X | -- | -- | 10 (100) | -- | -- | 9 (100) | -- | -- | 6 (100) |
| **(Total)** | **(Total)** | | | **(Total)** | | | **(Total)** | | |
| +/X x +/X | 8 (24.2) | 16 (48.5) | 9 (27.3) | 8 (25.8) | 14 (45.2) | 9 (29.0) | 11 (24.2) | 23 (48.5) | 13 (27.3) |
| +/X x X/X | -- | 14 (51.9) | 13 (48.1) | -- | 14 (50.0) | 14 (50.0) | -- | 16 (55.2) | 13 (44.8) |
| X/X x X/X | -- | -- | 27 (100) | -- | -- | 30 (100) | -- | -- | 24 (100) |

** Percentage of genotypes shown in parentheses ( ).*

**Supplemental Table 1. Genotype and survival distribution of *$Gdf11^{Gdf8aa}$*, *$Gdf8^{Gdf11aa}$*, and *$Gdf8^{Gdf11MD}$* F5 progeny in C57BL/6J background through three initial rounds of breeding.** Progeny of *$Gdf11^{Gdf8aa}$*, *$Gdf8^{Gdf11aa}$*, and *$Gdf8^{Gdf11MD}$* crosses were genotyped and analyzed to confirm Mendelian ratios of F5 progeny and rule out potential lethality as a result of the designed genetic modifications.
